# Structure and Function of TM6SF1 Reveal Role in mTORC1 Signaling

**DOI:** 10.64898/2026.04.01.715928

**Authors:** Sen Hong, Liangjie Jia, Rong Wang, Nadia Elghobashi-Meinhardt, Helen H. Hobbs, Xiaochun Li

## Abstract

The Transmembrane 6 Superfamily (TM6SF) comprises two members: TM6SF1, a ubiquitously expressed lysosomal membrane protein of unknown function, and TM6SF2, an endoplasmic reticulum protein required for bulk lipidation of Apolipoprotein B-containing lipoproteins. Here we used cryo-electron microscopy (cryo-EM) to determine the structure of human TM6SF1 at 2.9- Å resolution. TM6SF1 forms a polytopic homodimer, with each protomer comprising ten transmembrane helices (TMs). TMs 1-6 form a pocket that accommodates a cholesterol molecule. Cell-based assays revealed that loss of TM6SF1 perturbs mTORC1 signaling, resulting in reduced phosphorylation of S6 kinase 1 and 4E-BP1 and constitutive activation of Transcription Factor EB (TFEB) and that cholesterol is required for these effects. Biochemical analyses support the model that TM6SF1 directly engages LAMTOR1, a component of Ragulator complex, in a cholesterol-dependent manner. Together, these findings identify TM6SF1 as a lysosomal cholesterol binding protein involved in regulating mTORC1 signaling.

**Significance Statement:** We determine the cryo-EM structure of TM6SF1 and show that it is a lysosomal membrane protein that associates with LAMTOR1, a component of Ragulator complex, in a cholesterol-dependent manner. Loss of TM6SF1 impairs mTORC1 signaling and promotes nuclear accumulation of TFEB, suggesting a role for TM6SF1 in lysosome-dependent nutrient sensing and signaling. Together, our findings suggest that TM6SF1 is involved in the regulation of mTORC1 activity and provide insight into the functional diversification of the TM6SF protein family.

## Introduction

Lysosomes are membrane-bound organelles that function as the primary degradative compartments of eukaryotic cells. They contain a broad repertoire of hydrolytic enzymes that operate optimally under acidic conditions maintained by the vacuolar-type H⁺-ATPase (V-ATPase) (1, 2). Traditionally regarded as the intracellular recycling center, lysosomes mediate turnover of macromolecules delivered through endocytosis, phagocytosis, and autophagy (3–6). Defects in lysosomal enzymes, transporters, or membrane proteins cause a spectrum of lysosomal storage disorders that manifest as neurodegenerative, metabolic, and immune diseases. Beyond their canonical degradative function, lysosomes are dynamic signaling hubs that integrate cellular metabolism, nutrient sensing, and stress responses. Central to this regulatory function is the control of mTORC1 (mechanistic target of rapamycin complex 1) signaling, which coordinates cell growth and energy homeostasis in response to nutrient availability (7–10).

mTORC1, comprising mTOR, Raptor, and mLST8, is recruited from the cytoplasm to cytosolic face of lysosomes in response to amino acids. The Raptor subunit of mTORC1 binds the Rag guanosine triphosphatase (GTPase)-Ragulator complex, thus anchoring mTORC1 to lysosomes (11). Both lysosome-associated and non-lysosomal pools of mTORC1 regulate distinct signaling pathways, with their outputs depending on the source of metabolic inputs (12). Over 100 *bona fide* lysosomal transmembrane proteins have been identified by quantitative proteomic analyses (13, 14), including TM6SF1, a polytopic lysosomal membrane protein with 10 transmembrane helices (TMs) (15, 16). TM6SF1 belongs to the TM6SF family, which consists of two members: TM6SF1 and TM6SF2. Our interest in this family arose from the discovery of an inactivation mutation in TM6SF2 that confers susceptibility to steatotic liver disease (17). TM6SF2 is expressed primarily in the small intestines and the liver, residing in the smooth endoplasmic reticulum (ER).(18) The protein is required for optimal lipidation of ApoB-containing lipoproteins prior to secretion into the circulation (17–19). In contrast to TM6SF2, the physiological function of TM6SF1 is not known. Whereas the expression of TM6SF2 appears to be restricted to vertebrates, TM6SF1 is an evolutionarily conserved ancient protein present in metazoans and choanoflagellates, has a physiological function that remains unclear. The two members of the TM6SF family exhibit distinct subcellular localizations, implying divergent biological roles despite their similar architectures.

Structurally, TM6SF1 comprises two tandemly arrayed transmembrane repeats, each adopting an EXPERA (EXPanded EBP superfamily) fold that resembles the sterol isomerase EBP (emopamil-binding protein) (20) and the σ2 receptor (21). To elucidate the biological function of TM6SF1, we generated the TM6SF1 knockout (KO) cells for functional studies and we determined the structure of the protein at 2.9-Å resolution. The structure, coupled with biochemical studies, revealed that TM6SF1 binds a cholesterol molecule within the TM domains, an interaction required for association with the Ragulator complex and recruitment of mTORC1 to lysosomes. Loss of TM6SF1 disrupts mTORC1-dependent S6 kinase 1 and 4E-BP1 phosphorylation as well as TFEB phosphorylation, leading to nuclear entry and constitutive activation of genes involved in lysosomal and autophagosomal function. Together, these findings identify TM6SF1 as a cholesterol-binding lysosomal membrane protein that is essential for selective mTORC1 signaling events.

## Results

### Structure determination of TM6SF1

The small size of TM6SF1 (∼42 kDa) and absence of distinct extra-membranous features made cryo-EM particles alignment challenging. To increase particle size and facilitate alignment, we fused a maltose-binding protein (MBP) cassette to the N-terminus of TM6SF1. We linked the C-terminal helix of MBP to the N-terminus of the first transmembrane helix (TM) of TM6SF1 via a helical linker (AEEEKRKAEEEKRK). To further enhance image alignment, an MBP binder, designed ankyrin-repeat protein off7 (DARPin off7) (22, 23), was fused to the C-terminus of TM6SF1 through a flexible GS linker (GGGGSGGGGS), serving as a fiducial marker (Fig. 1*A*).

**Fig. 1.**
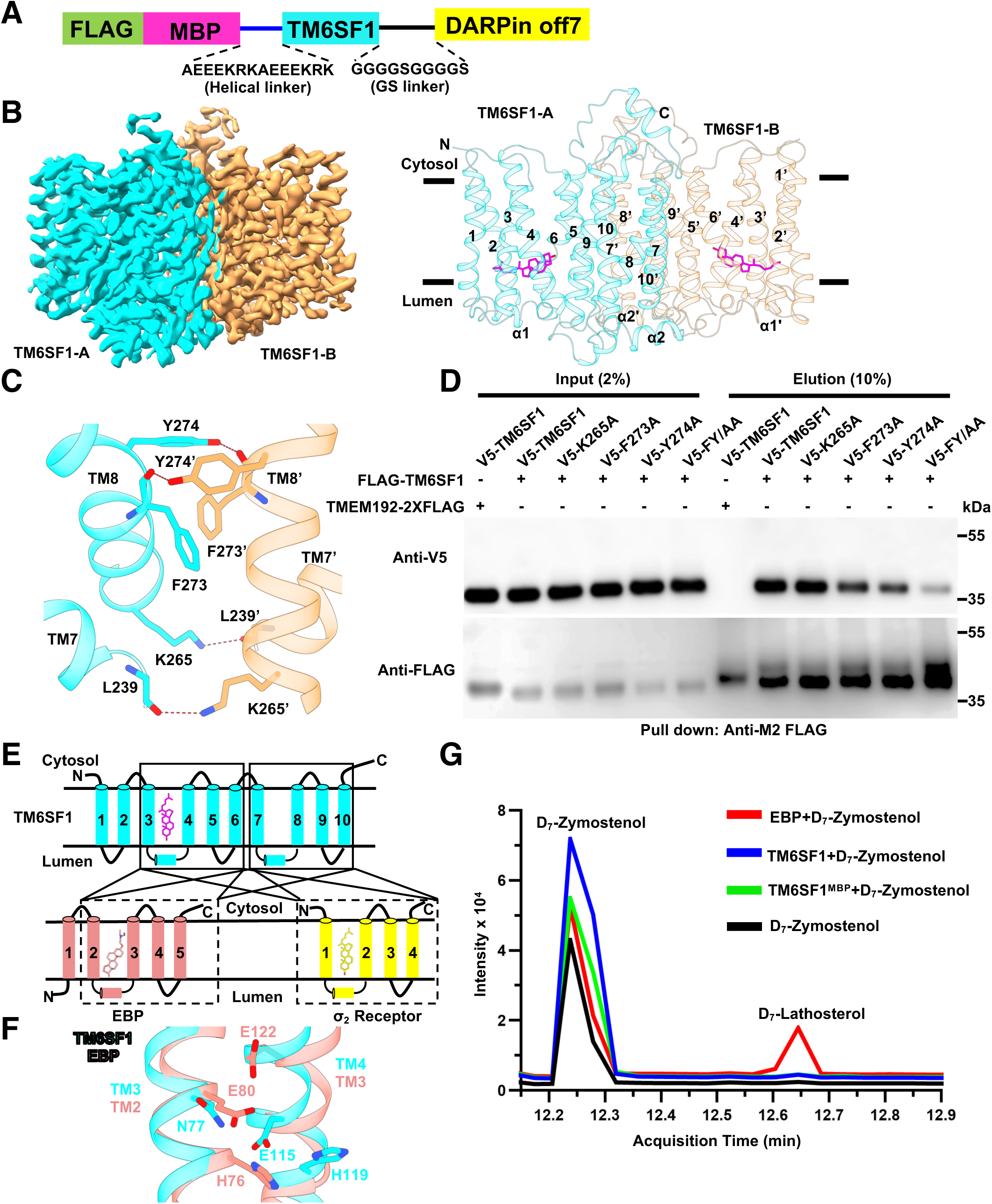
Structure of cholesterol-bound human TM6SF1. (*A*) Human TM6SF1^MBP^ expression construct for cryo-EM study. (*B*) Cryo-EM density map and overall structure of TM6SF1^MBP^. TM6SF1 forms a dimer, with the two monomers colored in cyan (TM6SF1-A) and brown (TM6SF1-B) (left), and with the TM helices numbered (right). (*C*) Dimeric interface of TM6SF1. Key interacting residues (F274, Y273, K265, L239) from each monomer are shown as sticks, with potential hydrogen bonds indicated by dashed lines. (*D*) Mutational analysis of the TM6SF1 dimer interface by Co-IP. Inputs and proteins eluded from anti-M2 FLAG resin were analyzed by immunoblotting with anti-V5 and anti-FLAG antibodies. (*E*) Schematic of shared structural features of TM6SF1 (cyan), EBP (lower left, pink) and σ2 receptor (lower right, yellow). *(F)* Structural alignment of TM6SF1 (cyan) with EBP (pink). A catalytic triad of EBP (His76, Glu80, and Glu122) forming the active core is shown as sticks. TM6SF1 contains similar residues in the corresponding positions, implying it may possess catalytic activity. (*G*) Gas Chromatography-Mass Spectrometry (GC-MS) analysis shows that EBP, but not TM6SF1, catalyzes conversion of zymostenol to lathosterol. D_7_-zymostenol was incubated alone (black) or together with EBP (red), TM6SF1 (blue), or TM6SF1^MBP^ (green), and the products were analyzed by GC-MS. EBP served as a positive control. The major peak at ∼12.25 min corresponds to the substrate D_7_-zymostenol. The product peak corresponding to D_7_-lathosterol (∼12.65 min) was detected only in reaction containing EBP.

This construct is henceforth referred to as TM6SF1^MBP^. TM6SF1^MBP^ was overexpressed in HEK293 GnTI⁻ cells and purified in n-Dodecyl-β-D-Maltopyranoside (DDM). For structural analysis, the detergent was replaced with digitonin during size exclusion chromatography (SEC) (*SI Appendix*, Fig. S1*A*). After 3D reconstruction with C1 symmetry, the cryo-EM map of TM6SF1 was clearly resolved (Fig. 1*B*), whereas the density of the second MBP-DARP complex was not well defined. To further enhance the density of TM6SF1, we applied a mask encompassing TM6SF1 and performed local refinement (*SI Appendix*, Fig. S1*B*). This approach yielded an overall resolution of 2.86 Å (*SI Appendix*, Fig. S1 *B-E* and Table S1). The secondary structure and the majority of the amino acid residues were clearly resolved in the cryo-EM map (*SI Appendix*, Fig. S2 *A* and *B*).

The cryo-EM structure shows TM6SF1 as a homodimer (TM6SF1-A and TM6SF1-B), with each protomer comprising 10 TM helices (Fig. 1*B*). Although no symmetry was imposed during cryo-EM data processing, TM6SF1-A and TM6SF1-B adopt nearly identical conformations, with a root-mean-square deviation (RMSD) of 0.306 Å (*SI Appendix*, Fig. S2*C*). The dimer interface between the TMs of TM6SF1-A and TM6SF1-B spans over 1000 Å^2^, with TM7 and TM8 contributing to dimer assembly (Fig. 1*C*). The side chain of Phe273 stabilizes the dimer interface via a π-π interaction with an interplanar distance of 3.8 Å. Additionally, the side chains of Lys265 and Tyr274, along with the main chains of Leu239 and Phe273, contribute to dimer stabilization through potential hydrogen bonds, with distances of 3.4 Å and 2.5 Å, respectively (Fig. 1*C*). Our co-precipitation assay showed that V5-tagged TM6SF1 associates with FLAG-tagged TM6SF1 when co-expressed in HEK293A cells (Fig. 1*D*). This interaction was reduced when we substituted two aromatic residues at the interface with alanine (TM6SF1^F273A^ and TM6SF1^Y274A^). An even greater reduction occurred when both residues were substituted with alanine (TM6SF1^F273A/Y274A^, named TM6SF1^FY/AA^).

Dali search (24) indicated that TM6SF1 shares structural features with EBP (emopamil-binding protein), also known as 3-β-hydroxysteroid-Δ8, Δ7-isomerase (20) and with the σ2 receptor (21). EBP encodes a cholesterol biosynthetic enzyme that converts zymostenol to lathosterol (25). The σ2 receptor is highly expressed in various cancer cell types and has been implicated in cholesterol trafficking (21, 26). Although no sequence identity is shared between TM6SF1 and either EBP or σ2 receptor, they all share so-called EXPERA (EXPanded EBP superfamily) domains. TMs 2-5 of EBP has a topology resembling TMs 3-6 and TMs 7-10 of TM6SF1 with RMSDs of 1.180 Å and 1.441 Å, respectively (Fig. 1*E* and *SI Appendix*, Fig. S3*A*). Likewise, TMs 1-4 of the σ2 receptor align with TMs 3-6 and TMs 7-10 of TM6SF1, showing RMSDs of 1.175 Å and 1.326 Å, respectively (Fig. 1*E* and *SI Appendix*, Fig. S3*B*). Within the EXPERA, EBP contains a conserved catalytic triad composed of His76, Glu80, and Glu122, while TM6SF1 possesses a corresponding catalytic core consisting of His119, Asn77, and Glu115 (Fig. 1*F*). Adjacent to the catalytic triad in EBP is a lipid binding domain, and this enzyme has been shown to catalyze conversion of zymostenol to lathosterol in the cholesterol biosynthetic pathway. Unlike EBP, incubation of purified TM6SF1 with zymostenol did not produce any detectable lathosterol by mass spectrometry (Fig. 1*G*).

### TM6SF1 binds cholesterol

A substrate-like density was seen in TM6SF1 that was surrounded by TMs 1-6; No similar substrate pocket was formed in TMs 7-10 (Fig. 2 *A* and *B*). Structural alignment of TM6SF1 with EBP revealed that the density corresponded to where EBP binds U18666A, an inhibitor of cholesterol biosynthesis and Niemann-Pick Type C1 (NPC1) (27) (Fig. 1*E* and *SI Appendix*, Fig. S3*A*). Furthermore, structural superposition of TMs 1-4 of the cholesterol-bound σ2 receptor with TMs 3-6 of TM6SF1 revealed that cholesterol occupies a similar position within the putative substrate pocket of TM6SF1 (Fig. 1*E* and *SI Appendix*, Fig. S3*B*). Given that all three structurally related proteins – TM6SF2,, EBP, and σ2 receptor – are involved in lipid metabolism, we speculated that the density in TM6SF1 was likely a lipid, and due to the shape of the pocket, most likely cholesterol (Fig. 2*B*). Therefore, we extracted lipids from purified TM6SF1 and TM6SF1^MBP^ and analyzed the lipids by Gas Chromatography-Mass Spectrometry (GC-MS). A peak corresponding to cholesterol was detected in the GC-MS profiles of purified TM6SF1^MBP^ (Fig. 2*C*). We performed a similar analysis with purified TM6SF1 to ensure that the retained cholesterol was not bound to MBP or DARPin off7, and the result was identical (Fig. 2*C*). We then performed an *in vitro* competitive assay. Deuterated cholesterol effectively competed with cold cholesterol, whereas 25-hydroxycholesterol did not (Fig. 2*D*).

**Fig. 2.**
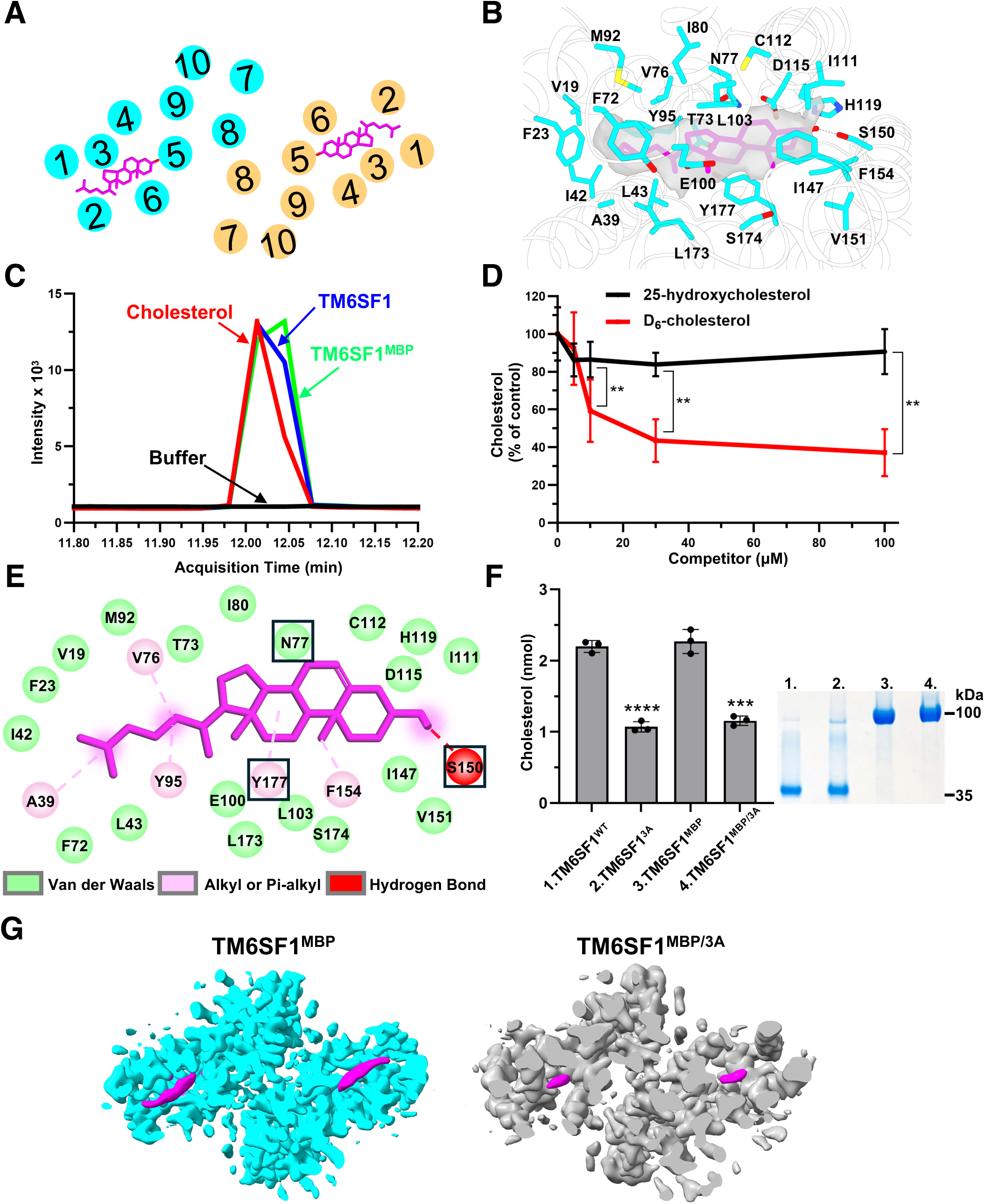
Cholesterol binding by TM6SF1. (*A*) Cross-sectional schematic of TM6SF1 showing the TM helices with location of cholesterol molecule (magenta sticks). (*B*) Details of molecular interactions between TM6SF1 and cholesterol. Cryo-EM map of cholesterol is shown in gray. (*C*) Chromatographs of sterols extracted from purified TM6SF1 using GC-MS. Chromatograms show the retention profiles of buffer (black), cholesterol (red), purified TM6SF1^WT^ (blue), and purified TM6SF1^MBP^ (green). The matching retention time between the cholesterol and sterol peaks in the TM6SF1 and TM6SF1^MBP^ samples confirms direct cholesterol binding to the protein. (*D*) D_6_-cholesterol competes with binding cholesterol to TM6SF1. Competitive binding assay measuring the displacement of cholesterol by increasing concentrations of D_6_-cholesterol (red) or 25-hydroxycholesterol (black). Data are presented as mean ± SE (n=3 independent experiments). Statistical significance was assessed by a two-tailed *t*-test. ** P < 0.01. (*E*) Key residues involved in Van der Waals (green), alkyl/Pi-alkyl (pink), and hydrogen bond (red) interactions are shown. Residue S150 forms a hydrogen bond with 3’-hydroxy group of cholesterol, suggesting its key role in substrate recognition. Residues mutated in the TM6SF1^3A^ are boxed. (*F*) Cholesterol quantification in purified TM6SF1^WT^, TM6SF1^3A^ and the corresponding cryo-EM constructs. Cholesterol extracted from 2 nmol of purified TM6SF1^WT^ or TM6SF1^3A^ or the corresponding cryo-EM constructs were quantified by GC-MS. Data represent mean ± SEM (n=3 independent experiments). ***p < 0.001, ****p < 0.0001 (two-tailed *t-test*). (*G*) Cryo-EM density maps of TM6SF1^MBP^ (cyan) and TM6SF1^MBP/3A^ (grey) displayed at the same contour level. To facilitate comparison, the TM6SF1^MBP^ map was low-pass filtered to the resolution of TM6SF1^MBP/3A^. Cholesterol-like densities (magenta) were clearly visible in the wild-type map but were markedly reduced in the mutant, consistent with impaired cholesterol binding upon mutation of the cholesterol-coordinating residues.

Structural analysis of the cholesterol binding pocket revealed that Ser150 in TM6SF1 binds the hydroxyl group of cholesterol, while several aromatic residues, including Phe23, Phe72, Tyr95, Phe154 and Tyr177, generate the hydrophobic environment to accommodate cholesterol (Fig. 2 *B* and *E*). We speculate that residues Val19, Phe23 and Ile42 may act as a gate to control access of cholesterol from the lipid bilayer to TM6SF1 (Fig. 2 *B* and *E*). To validate that TM6SF1 binds cholesterol, we substituted residues predicted to coordinate sterol binding to TM6SF1 with alanine to generate a triple mutant in both TM6SF1 and TM6SF1^MBP^ (N77A/S150A/Y177A, referred to as TM6SF1^3A^ and TM6SF1^MBP/3A^, respectively). Quantification of cholesterol binding to purified TM6SF1^WT^ and TM6SF1^3A^, as well as TM6SF1^MBP^ and TM6SF2^MBP/3A^, showed that the amount of cholesterol that bound the mutant proteins was ∼50% of WT TM6SF1. The residual cholesterol signal in the mutant proteins likely reflects cholesterol molecules associated with the transmembrane regions (Fig. 2*F*). We determined the cryo-EM structure of TM6SF1^MBP/3A^, yielding an overall resolution of 3.37 Å (*SI Appendix*, Fig. S4, Fig. S5 *A* and *B* and Table S1). The protein adopted a conformation that was almost identical to TM6SF1^MBP^, with a RMSD of 0.523 Å, except for the α1 helix, which protruded outward (*SI Appendix*, Fig. S5*C*). These data imply that the α1 helix of TM6SF1 may be involved in cholesterol binding. Comparison of the cryo-EM map of TM6SF1^MBP/3A^ to that of the wild-type TM6SF1^MBP^ at the same contour level revealed cholesterol-like densities in the wild-type protein that were reduced in the mutant (Fig. 2*G*). Together, these structural and biochemical data support assignment of the bound lipid density as cholesterol in the cryo-EM structure.

### TM6SF1 regulates mTORC1 signaling and transcription factor EB (TFEB) localization

Previously TM6SF1 was localized to lysosomes and shown to be expressed in the cerebellum, kidney, and intestine (16). To confirm its subcellular localization, we performed confocal microscopy of a fusion protein of TM6SF1 with GFP or mCherry in human HEK293A cells. The recombinant proteins colocalized with the lysosomal marker LAMP1, but not with markers of other organelles (calreticulin, ER; ERGIC, ER-Golgi intermediate complex; giantin, Golgi) (*SI Appendix*, Fig. S6). Then, we used CRISPR/CAS9 (28) to generate two TM6SF1 KO cell lines (KO-1 and KO-2) (Fig. 3*A*), as determined by genomic DNA sequencing. Neither cell line produced detectable TM6SF1 by immunoblotting using an anti-human TM6SF1 rabbit polyclonal antibody that we generated using a peptide from the C-terminus of TM6SF1 (Fig. 3*A*).

**Fig. 3.**
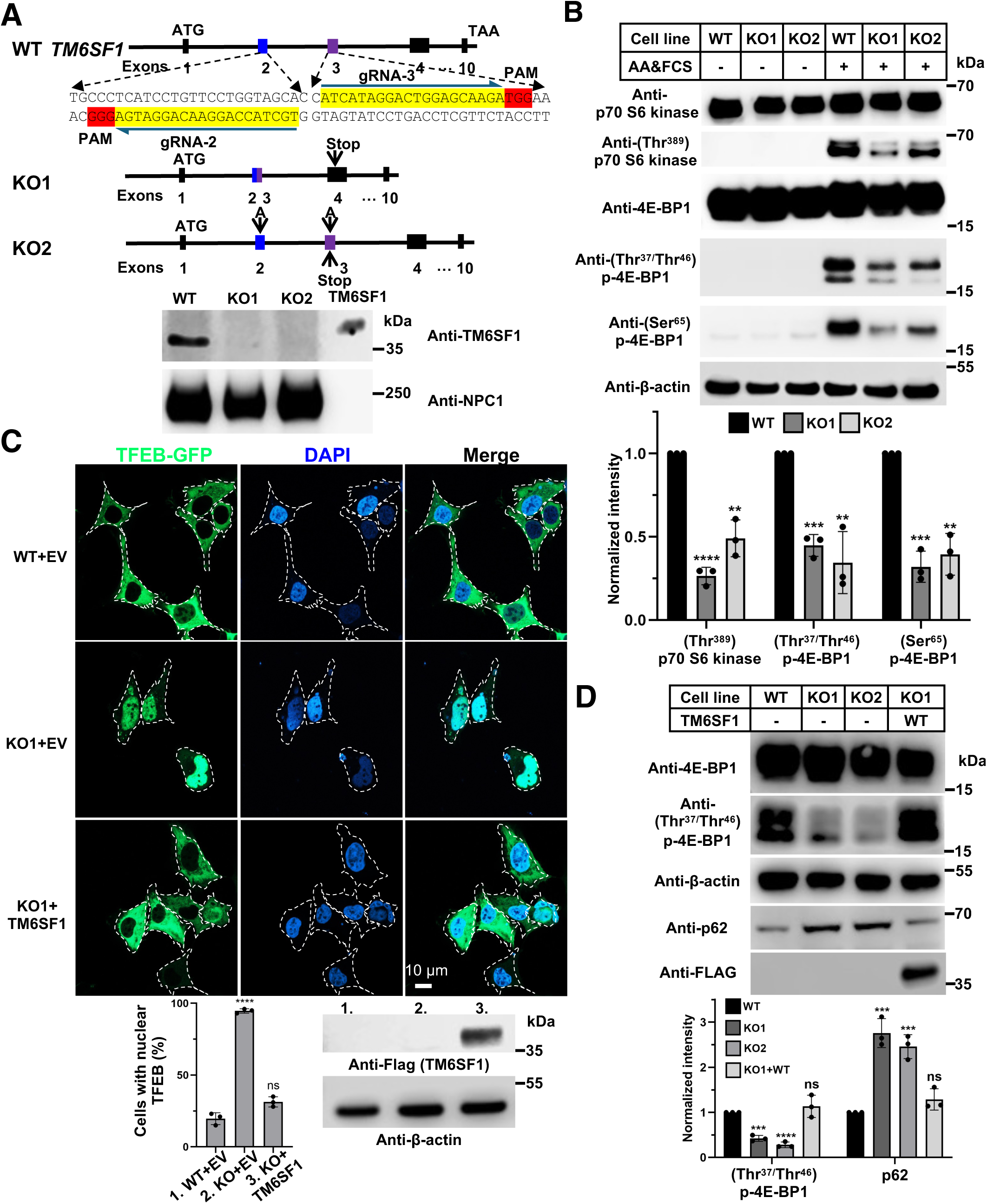
TM6SF1 contributes to mTORC1 signaling and TFEB localization. (*A*) CRISPR/Cas9-mediated generation of TM6SF1 knockout (KO) HEK293A cells. Two guide RNAs (gRNA2 and gRNA3) targeting exon 2 and exon 3 of human TM6SF1 are shown with target sequences highlighted in yellow and the PAM sequences in red. The guide RNAs were designed to induce deletion or insertion of the intervening region, leading to a frameshift and a premature stop codon in exon 4 (KO1) or exon 3 (KO2), respectively. Membrane fractions (10 μg) from knockout cells were confirmed to be deficient in TM6SF1 by immunoblotting with an anti-TM6SF1 antibody. NPC1 was used as a loading control. (*B*) Representative immunoblot analysis of mTORC1 pathway in TM6SF1 KO cells (KO1 and KO2 cells). On day 0, WT and TM6SF1 KO cells were set up in medium B supplemented with 10% FCS at a density of 1.5 × 10^5^ cells per well. On day 2, cells were incubated in HBSS for 1 hour to induce amino acid starvation. One group of cells was harvested immediately for immunoblotting, while the remaining cells were switched to medium B containing 10% FCS to allow amino acid refeeding for an additional 30 min before harvesting for immunoblotting. Phosphorylation levels of p70 S6 kinase (Thr³⁸⁹), 4E-BP1 (Ser^65^), and 4E-BP1 (Thr^37^/Thr^46^) were significantly reduced in KO cells, indicating attenuated mTORC1 signaling. β-actin was used as a loading control. Quantification of mTORC1 signaling is shown with mean ± SEM (n=3 independent experiments). Statistical significance was assessed by a two-tailed *t*-test with Welch’s correction. ** P < 0.01, *** P < 0.001, **** P < 0.0001. (*C*) TM6SF1 regulates TFEB nuclear localization. On day 0, WT and TM6SF1 KO cells were seeded on glass coverslips in medium B supplemented with 10% FCS. On day 1, cells were co-transfected with a TFEB plasmid and either an empty vector (EV), or FLAG-tagged human TM6SF1^WT^. On day 2, cells were then fixed and imaged using the confocal microscope. Representative confocal images show TFEB-GFP (green) localization in WT and KO cells. Nuclei are stained with DAPI (4’6-diamidino-2-phenylindole) (blue). Dashed lines indicate cell boundaries. Scale bar: 10 µm. Quantification of nuclear TFEB localization is shown below. Immunoblot analysis of TM6SF1 expression is also shown with β-actin used as a loading control. For each biological replicate, 100 cells were randomly observed and analyzed for nuclear TFEB localization. Data are presented as mean ± SEM (n=3 independent experiments). Statistical significance was assessed by two-tailed *t*-test or one-way ANOVA followed by Tukey’s multiple comparisons. ns, not significant; **** P < 0.0001. (*D*) Representative immunoblot analysis of p62 expression in TM6SF1 KO cells. Phosphorylation levels of 4E-BP1 (Thr^37^/Thr^46^) were reduced and expression levels of p62 increased in TM6SF1 KO cells. β-actin was used as a loading control. n=3 independent experiments. *** P < 0.001, **** P < 0.0001 (two-tailed *t-test*).

To determine if TM6SF1 participates in mTORC1 activation, we measured the phosphorylation status of selected mTORC1 target substrates in cells depleted and then replenished with amino acids. TM6SF1 deficiency attenuated mTORC1 signaling, as evidenced by decreased phosphorylation of p70 S6-kinase (Thr^389^) and 4E-binding protein 1 (4E-BP1) at Ser^65^ and Thr^37^/Thr^46^ (Fig. 3*B*), respectively. As an additional measurement of mTORC1 activity, we examined the subcellular distribution of GFP-tagged TFEB, a transcription factor that changes its cellular localization in response to phosphorylation by mTORC1. After mTORC1 is activated and recruited to lysosomes, it phosphorylates TFEB, so the protein cannot enter the nucleus and remains in the cytoplasm (29). If mTORC1 fails to be recruited to lysosomes or is inactivated, TFEB remains dephosphorylated and translocates into the nucleus where it activates genes involved in autophagy and lysosomal function, so-called CLEAR (Coordinated Lysosomal Expression and Regulation) genes (30)(31). In WT cells, TFEB was predominantly located in the cytosol, whereas in KO cells, it accumulated in the nucleus. The cytosolic localization of TFEB was rescued by expression of recombinant TM6SF1^WT^ in the KO cells (Fig. 3*C*). As anticipated based on this result, TM6SF1 KO cells had an increased level of a TFEB target proteins, p62 (SQSTM1) (32) (Fig. 3*D*). These data support a model in which TM6SF1 is required for mTORC1 phosphorylation of TFEB and its retention in the cytoplasm in response to extracellular stimuli.

We evaluated the effect of TM6SF1 deficiency on selected additional mTOR regulated lysosomal pathways. We monitored lysosomal acidification in WT and TM6SF1 KO1 live cells using two small-molecule fluorescent probes that accumulate in lysosomes: pHLys Red (pH-sensitive) and LysoPrime Green (pH-insensitive). The fluorescence intensity of pHLys Red increases with decreasing pH, whereas LysoPrime Green remains stable, irrespective of pH. No differences in color pattern were appreciated between the KO and WT cells, whereas treatment with bafilomycin A1 (33), a V-ATPase inhibitor, was associated with a marked increase in lysosomal pH, as expected (*SI Appendix*, Fig. S7).

We also examined whether TM6SF1 deficiency interferes with the transfer of cholesterol delivered to lysosomes by endocytosis across the lysosomal membrane, as occurs with inactivation of NPC1 (34). We compared the distribution of free cholesterol in the lysosomes of WT and KO cells by staining with filipin. No difference in the amount or distribution of filipin staining wase seen between WT and KO cells, whereas treatment with U18666A (27), an inhibitor of NPC1 caused a marked increase in filipin staining in lysosomes (*SI Appendix*, Fig. S8). Notably, lysosomes seemed larger and more dispersed in U18666A-treated KO cells than in WT cells. Together, these findings indicate that TM6SF1 does not cause a general disruption of lysosomal function but rather has a more defined effect on lysosomal mTOR signaling.

### TM6SF1 interacts with LAMTOR1

Next, we tested if the observed failure of mTORC1 signaling in selected pathways of the TM6SF1 KO cells was caused by changes in its interaction with other components of the mTORC1 lysosomal recruitment and signaling pathway. To identify such proteins, we immunoprecipitated TM6SF1 from HEK293A cells expressing GFP-tagged TM6SF1 and immunoblotted for selected lysosomal proteins. We found no evidence supporting a physical interaction between TM6SF1 and endogenous V1-ATPase (subunit A1) or SLC38A9, an arginine-sensitive lysosomal transmembrane protein involved in amino acid sensing (35, 36), whereas endogenous LAMTOR1, a component of the Regulator complex that links the mTORC1 complex to the lysosomal membrane co-immunoprecipitated with TM6SF1 (Fig. 4A)(37). To further confirm the physical interaction between TM6SF1 with LAMTOR1, we expressed and purified the Ragulator complex (LAMTOR 1-5) in *E.coli* as described (38) (Fig. 4*B*, bottom). The LAMTOR1 in the mixture had a hemagglutinin (HA) tag at the N-terminus. We then incubated the proteins in the presence and absence of FLAG-tagged TM6SF1. LAMTOR1 was pulled down by FLAG-TM6SF1 and conversely, TM6SF1 was pulled down by LAMTOR1, confirming a reciprocal interaction between the two purified proteins (Fig. 4*B*, top). We then performed gel filtration before (top gels) and after (bottom gel) mixing Regulator with TM6SF1 (2:1 ratio). The elution volume of TM6SF1 shifted from a peak between 16.5-17 mL (Fig. 4*C*, black) to 16-16.5 mL (Fig. 4*C*, blue) after the protein was mixed with the Ragulator complex, consistent with formation of a TM6SF1-Ragulator complex *in vitro* (Fig. 4*C*).

**Fig. 4.**
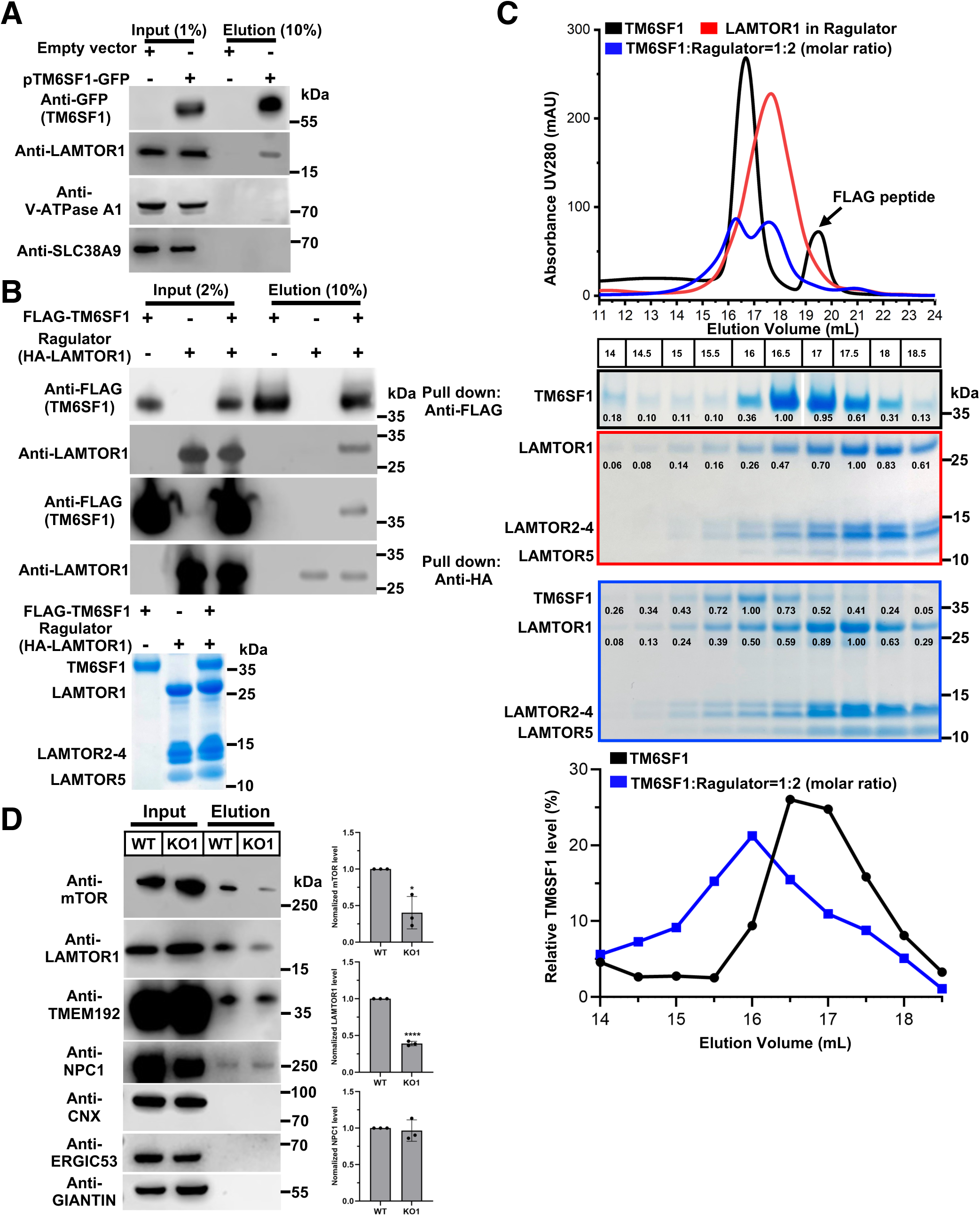
TM6SF1 interacts with LAMTOR1 of Ragulator complex. (*A*) TM6SF1 interacts with LAMTOR1. HEK293A cells were transfected with empty vector (EV) or GFP-tagged TM6SF1 (pTM6SF1-GFP). Proteins were immunoprecipitated from cell lysates using μMACS GFP microbeads. Immunoblot analysis was performed with antibodies against GFP, LAMTOR1, ATP6V1A, and SLC38A9. (*B*) Pull-down analysis validates interaction between TM6SF1 and LAMTOR1 *in vitro*. Coomassie-stained SDS-PAGE gel showing purified protein of TM6SF1 and Ragulator complex (left). FLAG-tagged TM6SF1 was immobilized on anti-FLAG M2 resin, while HA-tagged LAMOTR1 was captured using anti-HA resin. LAMTOR1, a component of the Ragulator complex, was expressed with an N-terminal HA tag and T7 tag connected by a 10-residue linker and migrated as a band of approximately 25 kDa on SDS-PAGE. Elution was analyzed by immunoblotting with anti-FLAG and anti-LAMTOR1 antibodies. (*C*) Gel filtration shows TM6SF1 bound to Ragulator complex in the presence of 0.06% Digitonin. Normalized GIANTIN, and ERGIC53 in whole-cell lysates and purified lysosomes. Quantification was performed using ImageJ. NPC1 was used as a control. Data represent mean ± SEM (n=3 independent experiments). *p < 0.05 (two-tailed *t-test*).

If the interaction between LAMTOR1 and TM6SF1 is required for recruitment of mTORC1 to lysosomes, we would expect that mTORC1 would not associate with lysosomes in the absence of TM6SF1. Intact lysosomes were selectively isolated from WT HEK293A cells and TM6SF1 KO cells using LysoTag (TMEM192-2xFLAG) (1), as evidenced by the presence of equal amounts of NPC1 and the absence of selected proteins from the ER (calnexin), ER-Golgi intermediate compartment (ERGIC53) and Golgi (giantin). The amount of mTOR and LAMPTOR associated with lysosomes was reduced in TM6SF1 KO cells compared to WT cells (Fig. 4D). These findings support a model in which TM6SF1 interacts with the Ragulator complex, reinforcing LAMTOR1-dependent membrane anchoring to establish a competent mTORC1 docking platform.

### Cholesterol modulates TM6SF1 in regulating mTOR signal

To determine whether cholesterol binding to TM6SF1 is required for mTORC1 docking and activation, we depleted cellular cholesterol in WT HEK293A and TM6SF1 KO cells and then replenished the cells with cholesterol for 2 hours, either providing the cholesterol to the cells in methyl-β-cyclodextrin (MCD), or by repleting the medium with low density lipoproteins (LDL) (Fig. 5*A*). In WT cells, cholesterol depletion markedly reduced phosphorylation of 4E-BP1 at Thr^37^/Thr^46^ and Ser^65^. Both cholesterol and LDL refeeding robustly restored these phosphorylation events. In contrast, cholesterol repletion had minimal effect on reactivating mTORC1 signaling in TM6SF1 KO cells (Fig. 5*A*). Importantly, expression of TM6SF1^WT^, but not cholesterol-binding defective TM6SF1^3A^, rescued 4E-BP1 phosphorylation in the KO cells refed with cholesterol or LDL (Fig. 5*A*). We documented that TM6SF1^3A^ localized to lysosomes in this experiment (SI Appendix, Fig. S9). These findings demonstrate that cholesterol binding to TM6SF1 is required for efficient mTORC1 activation.

**Fig. 5.**
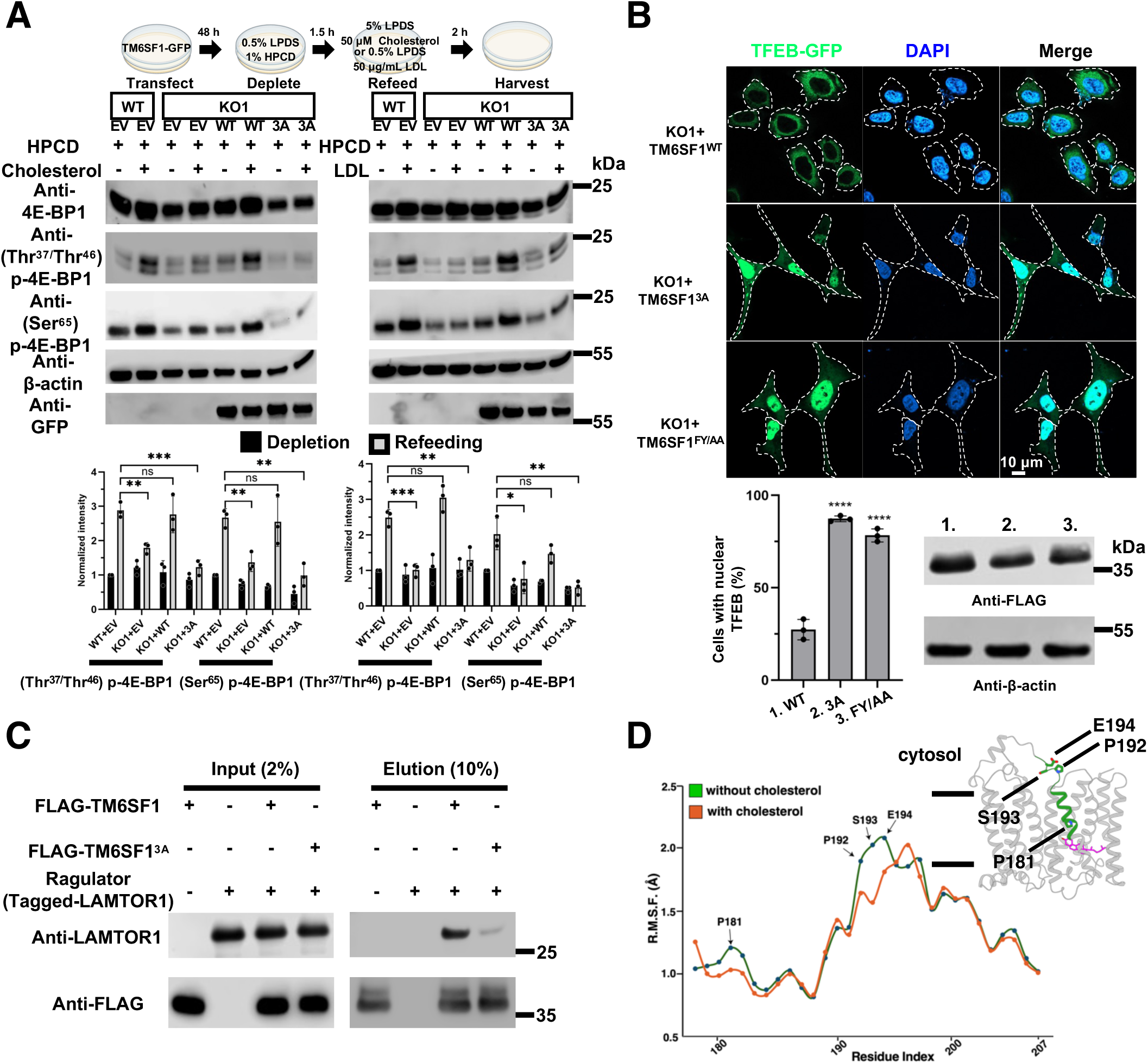
TM6SF1-mediated mTORC1 docking depends on cholesterol. (*A*) TM6SF1 is required for mTORC1 activation by cholesterol. TM6SF1 WT or TM6SF1 KO HEK293A cells were depleted of sterols using 1% HPCD for 1.5 hour, followed by refeeding 50 μM MCD-cholesterol (left) or 50 μg/mL LDL (right) for 2 hours. Cell lysates were immunoblotted with the indicated antibodies. Quantifications are performed using ImageJ and presented as mean ± SEM (n=3 independent experiments). Statistical significance was determined using a two-tailed Student’s *t-test* assuming equal variances. * P < 0.05, ** P < 0.01, *** P < 0.001. (*B*) Disrupting cholesterol binding (TM6SF1^3A^) or dimerization (TM6SF1^FY/AA^) affects TFEB nuclear localization. Representative confocal images show TFEB-GFP (green) localization in WT and KO cells. Nuclei are stained with DAPI (blue). Dashed lines indicate cell boundaries. Scale bar: 10 µm. Quantification of nuclear TFEB localization is shown at the bottom. Immunoblot analysis of TM6SF1 expression levels is also shown below. For each biological replicate, 100 cells were randomly analyzed for total and nuclear TFEB localization. Data are presented as mean ± SEM (n=3 independent experiments). Statistical significance was assessed by a one-way ANOVA followed by Tukey’s multiple comparisons. ****P < 0.0001. (*C*) Disrupting cholesterol binding attenuates the interaction between TM6SF1 and Ragulator complex. FLAG-tagged TM6SF1 was bound to anti-FLAG M2 resin and pulldown analysis was performed in buffer B. The input and elution were immunoblotted with the indicated antibodies. (*D*) MD simulations indicate the dynamics of residues 178-207 of TM6SF1 in the presence (orange) or absence (green) of cholesterol in the central pocket. Residues 179-194 in the structure are colored green.

To further study the requirement of cholesterol for TM6SF1-mediated mTORC1 signaling, we compared the pattern of TFEB localization in TM6SF1 KO cells expressing TM6SF1^WT^, TM6SF1^3A^, and TM6SF1^FY/AA^. Expression of TM6SF1^WT^ efficiently restored cytosolic TFEB localization (Fig. 5*B*). In contrast, expression of TM6SF1^3A^ or TM6SF1^FY/AA^ failed to rescue TFEB localization from the nucleus to the cytoplasm. These results indicate that cholesterol binding of TM6SF1 is required for TFEB relocalization in response to cholesterol, irrespective of whether the cholesterol is from the cell membrane or from the lysosomal lumen by endocytosis.

To assess whether cholesterol is required for TM6SF1 interaction with LAMTOR1, we performed pull-down assays using TM6SF1^WT^ and TM6SF1^3A^. Compared with TM6SF1^WT^, the interaction between TM6SF1^3A^ and LAMTOR1 was markedly reduced (Fig. 5*C*), suggesting that cholesterol binding is important for this interaction. We cannot rule out that the substituted sequences had effects on protein structure that were below our detection limits using cryo-EM (Fig. 2*G*). Finally, we performed molecular dynamics (MD) simulations to assess how cholesterol binding modulates the conformational dynamics of TM6SF1. In the cholesterol-free state, residues within TM6 (179–183), which lie proximal to the sterol-binding pocket, exhibited flexibility. While the adjacent cytosolic loop (192–194) showed an approximately 0.5 Å increase in Root Mean Square Fluctuation (RMSF) over 200-ns simulations compared with the cholesterol-bound condition (Fig. 5*D*). These results indicate that cholesterol might act as a structural brace within TM6SF1, restricting conformational flexibility and maintaining the cytosolic region in a geometry competent for LAMTOR1 engagement. The precise molecular determinants governing the interaction between TM6SF1 and LAMTOR1 remain to be elucidated.

## Discussion

Our study identifies TM6SF1 as a cholesterol-binding polytopic protein in the lysosomal membrane that contributes to mTORC1 signaling. The cryo-EM structure reveals a cholesterol molecule bound within a pocket formed by the TMs of TM6SF1 near the lysosomal luminal leaflet. Biochemical analyses further show that TM6SF1 associates with LAMTOR1, a membrane-anchoring subunit of the Ragulator complex. Disruption of the cholesterol-binding site in TM6SF1 weakens the physical interaction between TM6SF1 and LAMTOR1 and is associated with impaired mTORC1 signaling and constitutive activation of TFEB.

Two major pools of mTORC1 have been described: a cytosolic fraction and a lysosome-associated fraction, with recruitment to the lysosomal membrane being critical for full mTORC1 activation (39). Previous studies have shown that the heteropentameric Ragulator complex is anchored to the lysosomal membrane through lipid modifications on LAMTOR1 (40, 41). Specifically, Gly2 and Cys3/Cys4 of LAMTOR1 are N-myristoylated and S-palmitoylated, respectively, and these modifications are essential for its membrane-scaffolding function (41). We showed that LAMTOR1 also contributes to its interaction with TM6SF1, potentially facilitating the recruitment of mTORC1 to lysosomes.

LDL-derived cholesterol is transported to and released into the luminal leaflet of the lysosomal membrane by NPC1 (34, 42). Cholesterol delivered through this pathway may enter the binding pocket of TM6SF1 and promote, stabilize, or allosterically regulate its association with the Ragulator complex. In TM6SF1, cholesterol is deeply embedded within the transmembrane region near the luminal leaflet, representing a binding mode distinct from that of LYCHOS, a GPCR-transporter chimeric protein proposed to sense lysosomal cholesterol and regulate GATOR1-dependent control of RagA/B (43) (*SI Appendix,* Fig. S10). Loss of LYCHOS reportedly does not impair mTORC1 signaling under amino acid-replete conditions but selectively attenuates mTORC1 activation upon cellular cholesterol depletion (44). In contrast, loss of TM6SF1 disrupts amino acid-stimulated mTORC1 activation. Although this comparison suggests that TM6SF1 and LYCHOS may affect lysosomal mTORC1 signaling through distinct mechanisms, any direct functional relationship between these two proteins remains speculative.

Several limitations of the present study merit explicit acknowledgment. First, all experiments were performed in HEK293A cells. Whether TM6SF1 performs an analogous scaffolding function in other cellular contexts, particularly in tissues with high lysosomal activity and cholesterol turnover such as brain, macrophages or neurons remains unclear. Second, the MBP-TM6SF1-DARPin off7 fusion construct was designed specifically as a structural tool to stabilize TM6SF1 for cryo-EM study. Because MBP and DARPin off7 assemble on the cytosolic face of TM6SF1, they may sterically alter the binding of LAMTOR1. This construct should therefore be viewed as a structural tool rather than a functional surrogate. For functional studies we did not use a fusion protein of TM6SF1. Third, TM6SF1-deficient cells may have more general abnormalities in lysosomal structure, raising the possibility that the observed defects in amino acid-stimulated mTORC1 signaling could be either a direct consequence of impaired Ragulator scaffolding or an indirect consequence of broader lysosomal dysfunction. Systematic evaluation of lysosomal integrity, including lysosomal protease activity, autophagic flux, membrane permeability, and other markers of lysosomal function, will be necessary to determine the selectivity of the defect.

Collectively, the structural and biochemical data presented here support a model in which cholesterol-bound TM6SF1 functions as a lysosomal membrane scaffold that interacts with LAMTOR1 and promotes mTORC1 recruitment to the lysosomal surface (*SI Appendix*, Fig. S9). This model places TM6SF1 within a broader framework in which lysosomal membrane cholesterol is coupled to mTORC1 activity, potentially through multiple parallel sensing or scaffolding mechanisms. Testing this model in additional cell types and defining whether the mTORC1 signaling defect reflects a direct impairment of Ragulator docking or an indirect consequence of altered lysosomal physiology will be important directions for future work.

## Acknowledgements

The Cryo-EM data were collected at the UTSW Cryo-EM Facility and HHMI Janelia Cryo-EM Facility. This research was supported in part by the computational resources provided by the BioHPC supercomputing facility located in the Lyda Hill Department of Bioinformatics, UT Southwestern Medical Center, TX. URL: https://portal.biohpc.swmed.edu. We would like to thank Lisa Kinch and Jonathan Cohen for helpful discussions, and Christina Zhao and Fang Xu for excellent technical support. We thank Lisa Beatty for cell culture work and Linda Donnelly for antibody generation. We thank Jeffrey McDonald of the Lipid Mass Spectrometry Core and Andrew Lemoff in the Proteomics Core, both at UTSW, for measurement of tissue lipid and protein quantification, respectively. This work was supported by NIH-R01DK090066, P30DK127984, P01-HL1600487, R35 GM149533, and Welch Foundation (I-1957).

## Author Contributions

S.H., H.H.H., and X.L. designed the experiments; S.H. determined the structures; S.H., J.L. and R.W. conducted the functional experiments; N.E.M. performed the MD simulations; all the authors analyzed data and S.H., H.H.H. and X.L. wrote the paper.

## Data, code, and materials availability

All data are available in the main text or the supplementary materials. The 3D cryo-EM density maps of TM6SF1^MBP^ and TM6SF1^MBP/3A^ have been deposited in the Electron Microscopy Data Bank (EMDB) under accession numbers EMD-75471 and EMD-75606, respectively. The corresponding atomic coordinates have been deposited in the Protein Data Bank (PDB) under accession numbers 10UP and 11BR, respectively.

