## Supplementary information for "Structure and Function of TM6SF1 Reveal Role in mTORC1 Signaling"

**This PDF file includes:**

SI Materials and Methods
SI Appendix, Figures S1 to S10
SI Table S1
SI References

### Methods

#### Protein expression and purification of human TM6SF1

The cDNA encoding human TM6SF1 (NM_023003.5) was subcloned into pEG BacMam (43) with a FLAG tag or V5 tag added to the N-terminus. For the TM6SF1$^{MBP}$ or TM6SF1$^{MBP/3A}$ construct, an N-terminal FLAG tag was placed upstream of the MBP coding sequence, followed by a helical linker, TM6SF1 or TM6SF1$^{3A}$, a glycine-serine (GS) linker, and a C-terminal DARPin off7 (22). A two-step PCR method was used to introduce mutations into the coding region of the *TM6SF1* cDNA. The coding sequences of the mutant constructs were validated by Sanger sequencing. Proteins were expressed in HEK293 GnTI$^-$ cells using the baculovirus system. Baculoviruses were generated by transfecting *Sf9* cells with Bacmid DNA using Cellfectin™ II Reagent. After two rounds of amplification, the resulting viral stocks were used to infect HEK293 GnTI$^-$ cells at a density of 4x10$^6$ cells/mL. After 8 hours at 37°C, sodium butyrate was added to a final concentration of 10 mM and the temperature was reduced to 30°C. After 72 hours, cells were homogenized in buffer A (20 mM HEPES, pH 7.5, 150 mM NaCl) supplemented with proteases inhibitor cocktail (Roche), leupeptin (10 μg/mL) and PMSF (1 mM) and then lysed by sonication. Cell debris was removed by low-speed centrifugation, and the supernatant was incubated with 1% (w/v) n-Dodecyl-β-D-Maltopyranoside (DDM) for 1 h at 4 °C. The insoluble fraction was removed by centrifugation (15,000 rpm, 4°C, 30 min) and the supernatant was loaded onto an anti-FLAG M2 resin (Sigma) that had been pre-equilibrated with buffer B (buffer A plus 0.02% DDM). The resin was washed once with 15 ml of Buffer B by gravity flow. The target protein was eluted in 7.5 mL of buffer C [buffer B supplemented with 3×FLAG peptide (0.1 mg/mL)] and initially purified by size-exclusion chromatography (SEC) using a Superose 6 Increase 10/300 GL column pre-equilibrated with buffer B. The purified samples were stored at -80 °C for future analysis. For cryo-EM sample preparation, peak fractions containing TM6SF1$^{MBP}$ or TM6SF1$^{MBP/3A}$ were concentrated and subjected to a second round of SEC using the same column, pre-equilibrated with buffer D (buffer A supplemented with 0.06% digitonin). The final peak fractions were concentrated and used for cryo-EM grid preparation.

#### Cryo-EM sample preparation and data collection

A total of 3 μL of purified protein at ~10 mg/mL was applied to glow-discharged Quantifoil R1.2/1.3 400 mesh Au holey carbon grids (Quantifoil). Grids were blotted using a Vitrobot Mark

IV (FEI) with a blotting force of 15, blotting time of 4 s, at 100% humidity at 22 °C and then plunged into liquid ethane cooled by liquid nitrogen. For TM6SF1$^{MBP}$, a total of 5,724 raw movie stacks were collected using SerialEM on a 300 kV Titan Krios (FEI) equipped with a Falcon4i detector. Data were acquired at a physical pixel size of 0.738 Å per pixel and a nominal magnification of 165,000. The defocus range was set to -0.8 ~ -1.8 μm. Each movie stack was recorded over 4 s and dose-fractionated into 40 frames with a total dose of ~ 60 electrons per A$^2$. For TM6SF1$^{MBP/3A}$, 3,087 raw movie stacks were recorded using a K3 camera at a 0.834 Å per pixel and a magnification of 105,000 with defocus range of -0.8 ~ -1.8 μm. Images were collected in super resolution mode with an exposure time of 5 s, dose-fractionated into 50 frames, and a total dose of ~ 60 electrons per A$^2$.

### Cryo-EM data processing

For TM6SF1$^{MBP}$ dataset, each raw movie stack (0.738 Å per pixel) was gain-normalized and patch motion correction and patch CTF estimation were performed using CryoSPARC v4 (44). Micrographs unsuitable for further analysis were removed through manual curation exposures. Subsequent processing steps, including blob-based particle auto-picking, 2D classifications, *ab initio* reconstruction, heterogeneous refinement, non-uniform refinement, and local refinement were conducted in CryoSPARC v4. Good classes from 2D classification were selected to generate three *ab initio* maps, which were then used for heterogeneous refinement. After two rounds of heterogeneous refinement, the best class was selected for non-uniform refinement. The resulting map was used to remove the detergent micelle and subsequently to generate masks in RELION 3.1.4 (45). The generated masks were used for particle subtraction to remove MBP and DARPin off7, resulting in the final map.

For TM6SF1$^{MBP/3A}$ dataset, raw movie stacks (0.834 Å per pixel) were also gain-normalized. Motion correction for beam-induced motion was performed using MotionCor2 (46) and the contrast transfer function (CTF) was estimated using CTFFIND4 (47) within Relion 3.1.4. A total of 618,456 particles were automatically picked using crYOLO-v1.9.7 with the general model and a particle threshold of 0.3 (48). Particles extractions were carried out in RELION3.1.4 using a box size of 350 pixels. Subsequent 2D classification, *ab initio* reconstruction, heterogeneous refinement, non-uniform refinement and local refinement was performed in CryoSPARC v4.

Masks used for local refinement were generated in RELION 3.1.4 and subsequently used for particle subtraction to remove MBP and DARPin (off7), resulting in the final map of the TM6SF1 and TM6SF1$^{(NSY/AAA)}$.

**Model building and refinement**

The initial model was built *de novo* by AlphaFold2 and docked into the cryo-EM map using ChimeraX (49). Manual model adjustments were performed in COOT(50), followed by real-space refinement in Phenix (51). Structural model validation was carried out using Phenix and MolProbity (52), with the results summarized in Extended Data Table 1. All structural figures were prepared using ChimeraX and PyMOL (www.pymol.org).

**Identification and quantification of sterols by GC-MS**

Sterols were extracted and analyzed following a modified saponification and derivatization protocol (53). Briefly, 15 µL of internal standard (ISTD; 0.67 mg/mL 5α-cholestane and 0.27 mg/mL epicoprostanol, equivalent to 5 µg and 4 µg, respectively) was added to 2 nmol purified protein. Then 1 mL of freshly prepared hydrolysis solution (6 mL of 10 mol/L KOH stock solution with 94 mL of ethanol) was added to the sample. Tubes were heated at 85°C for 2 hours for saponification with vortexing. After cooling to room temperature, 1 mL of PBS was added, and tubes were vortexed.

Sterols were extracted twice by adding 2 mL of petroleum ether to each tube, vortexing vigorously for 10-15 sec, and centrifuging at 2,800 rpm for 5 min. The upper petroleum phases from both extractions were combined in a clean test tube and mixed thoroughly. The combined organic phase was then dried under a stream of nitrogen gas. Next, 100 µL of Tri-Sil reagent (derivatization reagent) was added to each dried extract and vortexed. The solution was transferred to a GC vial and incubated at 75°C for 15 min, then cooled to room temperature. Samples were transferred to a GC insert, which was placed back into a GC vial for analysis.

Samples were analyzed using an Agilent 7890B GC/5977A MSD by EI equipped with a Rxi-5Sil MS column (40 m × 0.180 mm ID; 0.18 µm film thickness) from RESTEK. The injection volume was 1 μL with a port temperature of 280 °C. Hydrogen was used as the carrier gas at a constant flow rate of 3.0 mL/min. The GC oven was initially held at 150°C for 2 min, ramped up to a temperature of 300°C at 30°C/min, and held for 16 min, yielding a total run time of 23 min. The sterols were analyzed in selected ion-monitoring mode, and the data was normalized to an internal standard prior to processing using Mass Hunter software (Agilent). All experiments were performed at least three times with similar results.

**[$^2$H]-Cholesterol competitive binding assays**

Each reaction containing 2 nmol purified proteins in 200 μL of buffer B with varying concentrations of competitor compounds (25-hydroxycholesterol or $D_6$-cholesterol). The mixtures were incubated for 1 hour at 4°C. After incubation, 40 μL of a 50% suspension of Anti-FLAG M2 Magnetic Beads (Sigma), pre-equilibrated with buffer B, was added to each reaction and incubated for an additional hour at 4°C. After incubation, the magnetic beads were washed three times with buffer B using a DynaMag-2 magnetic rack (Invitrogen). FLAG-tagged proteins were eluted with 200 μL of buffer C. The eluted protein was subsequently analyzed for sterol using gas chromatography-mass spectrometry (GC-MS). This experiment was repeated twice and the results were similar.

**Enzymatic activity analysis in vitro**

A total of 2 nmol of purified protein in buffer B was incubated with 50 μM $D_7$-zymostenol (Avanti) at 37°C for 12 hours in a final reaction volume of 200 μL. Purified EBP protein (20) was used as a positive control, while buffer B alone served as the negative control. The enzymatic reactions were quenched by 10 μL of chloroform/methanol (2:1, v/v). Subsequent steps for sterol extraction, derivatization, and GC-MS analysis followed the same procedure described above for sterol identification and quantification. Substrate and product were analyzed in selected ion-monitoring (SIM) mode. This experiment was repeated twice and the results were similar.

**Cell culture and plating conditions**

Medium A is Opti-MEM$^{TM}$ I Reduced Serum Medium (Opti-MEM medium, Gibco). Medium B is Dulbecco's modified Eagle's medium (DMEM)-high glucose (4500 mg/L). Medium C is a 1:1 mixture of Ham's F-12 medium and DMEM. The cholesterol-depletion medium consists of medium C supplemented with 5% newborn lipoprotein-deficient serum (LPDS), 30 µM compactin, and 200 µM mevalonate (CPN/MEV). All mediums are supplemented with 100 U/mL of penicillin and 100 µg/mL of streptomycin sulfate. Cells were cultured under standard conditions at 37 °C in a humidified incubator with 8.8% $CO_2$.

**Generation of polyclonal rabbit anti-human TM6SF1 antibodies**

A polyclonal rabbit anti-TM6SF1 antibody (673E) was generated using keyhole limpet hemocyanin (KLH)-conjugated peptides on cysteine corresponding to the C-terminal 18 amino acids of human TM6SF1 (CIYKPEFFIKTKAEEKVE). Immunoblot shows that the 673E recognizes human TM6SF1.

**Generation of TM6SF1 KO cell lines**

CRISPR-Cas9 technology (28) was used to inactivate TM6SF1 gene in HEK293A cells. Briefly, two guide RNAs [gRNA2: TGCTACCAGGAACAGGATGA (5' → 3') and gRNA3: ATCATAGGACTGGAGCAAGA (5' → 3')], targeting exon 2 and exon 3 of the human TM6SF1 gene, respectively, were designed using the Benchling CRISPR design tool (http://www.benchling.com/crispr). The guide oligonucleotide pairs were annealed and cloned into pX459 v2.0 plasmid and then used to transfect HEK293A cells. Cells were transfected with 1 µg of each gRNA plasmid using FuGENE 6 (Promega) at a 5:1 FuGENE 6: DNA ratio in medium A. After 4 hours, medium A was supplemented with 10% fetal calf serum (FCS), and cells were incubated for an additional 48 hours. Cells were then trypsinized and subjected to puromycin selection (1 µg/mL) in medium B supplemented with 10 % FCS. After 10-14 days, Single-cell cloning was performed by limiting dilution (2 cells/well) in 96-well plates. Surviving cells were expanded into 24-well plates and further propagated. Membrane fractions from single clones were screened for TM6SF1 expression by immunoblotting. NPC1 (ab134113, Abcam) was used as a loading control. Clones that lacked TM6SF1 expression after at least three passages were designated TM6SF1 KO cell lines.

**mTORC1 signaling assay**

Detailed incubation conditions are provided in the figure legends. Amino acid (AA) starvation assays were performed as described previously (54). Briefly, cells were washed twice with pre-warmed HBSS containing calcium and magnesium and then incubated in the same buffer for 1 h to fully inhibit mTORC1 activity. For amino acid refeeding (+AA), HBSS was aspirated and cells were incubated in medium B supplemented with 10% FCS for an additional 30 min to achieve maximal activation of mTORC1 before harvest for Western blotting. Cells were lysed in RIPA buffer supplemented with a protease inhibitor cocktail (Roche), 100U/mL Benzonase Nuclease (Sigma), and PhosSTOP™ (Roche) phosphatase inhibitor at 4℃ for 20 min. Cell debris was removed by high-speed centrifugation at 16,100 x *g* for 10 min at 4°C, and the supernatant was collected. Protein concentrations were determined using the BCA assay (Thermo Fisher Scientific) and equal amounts were mixed with 4x SDS loading dye. The proteins were separated on 4-12% gradient SDS-PAGE gels and analyzed by western blotting with the indicated antibodies. This experiment was repeated at least twice and the results were similar.

Primary antibodies used were anti-p70 S6 kinase (9202S, Cell Signaling Technology (CST)), anti-Phospho-p70 S6 kinase (Thr389) (9234S, CST), anti-4E-BP1 (9452S, CST), anti-Phospho-4E-BP1 ($Ser^{65}$) (9451S, CST), anti-Phospho-4E-BP1 ($Thr^{37}$/$Thr^{46}$) (2855S, CST), anti-P62 (95697S, CST), anti-β-actin (4967S, CST). The secondary antibody used was HRP-conjugated goat anti-rabbit IgG (H+L) (111-035-144, Jackson ImmunoResearch).

**Immunofluorescence analysis**

For TFEB-EGFP imaging, incubation details are provided in the figure legends. Cells were fixed in 4% paraformaldehyde (Electron Microscopy Sciences) for 15 min at room temperature, washed three times with PBS, and mounted using Prolong Gold antifade reagent with DAPI (4'6-diamidino-2-phenylindole) (Invitrogen). Images were obtained using a Zeiss LSM800 confocal microscope. All images within the same figure panel were acquired using the same software setting. This experiment was repeated twice and the results were similar.

#### Lysosomal pH assay

Detailed incubation conditions are provided in the figure legends. According to the manufacturer's protocol, cells were washed twice with HBSS and incubated with 200 μL of LysoPrime Green working solution (1:2000 dilution) at 37°C for 30 min. After incubation, the dye was discarded, and cells were washed twice with HBSS before the cells were incubated at 37 °C for 30 min in 200 μL of pHLys Red working solution (1:1000 dilution in HBSS) in the presence or absence of bafilomycin A1 (1:1000 dilution), was added to the cells and incubated. The solution was then removed, and cells washed twice with HBSS. Medium B supplemented with 10 % FCS was added to the wells, and the stained cells were mounted using Prolong™ Gold antifade reagent (Invitrogen). Imaging was acquired using a Zeiss LSM800 confocal microscope. All images were performed with the same acquisition settings. This experiment was repeated twice and the results were similar.

#### Visualization of cholesterol distribution by fluorescence microscopy

Detailed incubation conditions are provided in the figure legends. Cells were fixed in 4% (w/v) formaldehyde in PBS for 30 min at room temperature, followed by three washes with PBS. Fixed cells were stained with 25 μg/mL filipin (Sigma) in PBS for 30 min at room temperature and washed three additional times with PBS. Coverslips were then mounted using Prolong™ Gold antifade reagent (Invitrogen). Fluorescence images were acquired using a Zeiss LSM800 confocal microscope equipped with a Plan-Apochromat 63×/1.4 oil DIC objective and ZEN imaging software (Zeiss, Oberkochen, Germany). All images of different conditions were captured with the same acquisition settings. This experiment was repeated twice and the results were similar.

#### Protein expression and purification of human Ragulator

The Ragulator complex was purified from *E.coli* BL21 cells (ThermoFisher), as previously described (36). Briefly, genes encoding Ragulator subunits were amplified from genomic DNA by PCR. LAMTOR1 was cloned into pET28a with an N-terminal His, HA tag, or both tags. LAMTOR2 and LAMTOR3 were inserted into pACYCDuet-1 without tags, whereas LAMTOR4 and LAMTOR5 were cloned into pETDuet-1 without tags. Plasmids encoding LAMTOR1-5 were co-transformed in *E.coli* BL21 cells, and protein expression was induced with 0.5 mM IPTG when the culture reached by OD600 of ~0.8, followed by incubation at 18 ℃ for approximately 12 h. Cells were harvested by centrifugation, resuspended in buffer A supplemented with 1 mM PMSF, and lysed by sonication. After removal of cell debris by centrifugation, the supernatant was applied to Ni-NTA resin pre-equilibrated with buffer A. The resin was washed with buffer A containing 25 mM imidazole, and the proteins were eluted with buffer A plus 250 mM imidazole. The eluted proteins were further purified by size-exclusion chromatography using a Superose 6 Increase 10/300 GL column pre-equilibrated with buffer A. Peak fractions containing the Ragulator complex were collected and stored at -80°C for subsequent analysis.

**Co-immunoprecipitation (Co-IP)**

To identify TM6SF1-interacting proteins, HEK293A cells were transfected with GFP-tagged TM6SF1 using FuGENE6 (Promega) or infected with P2 viral stocks expressing GFP-tagged TM6SF1. Cells transfected or infected with empty vector served as negative control. After 24 h incubation, cells were harvested and lysed in cold lysis buffer (50 mM Tris, 150 mM NaCl, 1% Triton X-100, pH 7.5) supplemented with proteases inhibitor cocktail (Roche) and 100 U/mL of Benzonase Nuclease for 30 mins on ice. Lysates were centrifuged at 1,000 × *g* for 10 min at 4°C, and protein concentrations were determined using the BCA assay. Co-immunoprecipitation was performed using commercial µMACS GFP Microbeads Isolation Kit (Miltenyi Biotec). Briefly, 50 µL of µMACS GFP Microbeads were mixed with 500 µg of protein for 1 hour on ice. The mixture was loaded onto µMACS separation columns pre-equilibrated with lysis buffer, then washed and eluted according to the manufacturer's protocol. Eluted proteins were resolved by SDS-PAGE and analyzed by western blotting with the indicated antibodies.

To validate the interaction between TM6SF1 or TM6SF1$^{3A}$ and LAMTOR1, Purified TM6SF1 or TM6SF1$^{3A}$ was incubated with either WT Ragulator or LAMTOR1-mutant Ragulator at a 1:1 ratio at 4 °C for 1 h. Subsequently, 20 µL of a 50% suspension of anti-FLAG M2 magnetic beads (Sigma) was added to the mixture and incubated for an additional 1 hour at 4°C. Beads were washed with buffer B and eluted with buffer C. Eluted proteins were separated by SDS-PAGE and analyzed by western blotting using indicated antibodies.

To validate the dimer interface of TM6SF1, HEK293A cells expressing WT or mutant TM6SF1 were infected with different P2 viral stocks. TMEM192 was used as a negative control. Cells were harvested using a cell scraper, rinsed with PBS and lysed in cold lysis buffer as described above. Lysates were centrifuged at 1,000 × g for 10 min at 4°C, and protein concentrations were determined using the BCA Assay Kit. A total of 20 µL of a 50% suspension anti-FLAG M2 beads (Sigma) was mixed with 150 µg of protein for 1 hour on ice. Beads were washed with buffer B and eluted with buffer C using a DynaMag-2 magnetic rack. Eluted proteins were separated by SDS-PAGE and immunoblotted with the indicated antibodies. Signal detection was performed using the SuperSignal$^{TM}$ West Pico PLUS (Thermo Scientific). Chemiluminescent signals were captured with the Odyssey FC Imager (LI-COR) and analyzed using Image Studio lite v5.2. The experiment was independently repeated twice on different days with similar results.

Primary antibodies used were anti-GFP (MA5-15256-HRP, Invitrogen), anti-V-ATPase A1 (sc-293336, Santa Cruz), anti-LAMTOR1 (8975S, CST), SLC38A9 (HPA043785, Sigma), anti-V5 (R960-25, Invitrogen), and anti-FLAG (F7425, Sigma). The secondary antibodies used were HRP-conjugated anti-mouse IgG antibody (7076S, CST), and HRP-conjugated goat anti-rabbit IgG (H+L) (111-035-144, Jackson ImmunoResearch).

**Gel-filtration analysis**

Purified TM6SF1 and Ragulator were mixed at a 1:2 molar ratio and then incubated on ice for 1 h. The mixture after the incubation was subjected to size-exclusion chromatography (SEC) using a Superose 6 Increase 10/300 GL column pre-equilibrated with buffer D. Eluted fractions were analyzed by SDS-PAGE (4-12% Bolt gels, Thermo Fisher Scientific) and visualized with InstantBlue Coomassie Stain (Abcam). This experiment was repeated twice and the results were similar.

**Immunopurification of lysosomes (LysoIP)**

Lysosomes were purified as described previously (1). Briefly, on day 0, WT and TM6SF1 KO cells were seeded in medium B supplemented with 10% FCS at a density of $1.5 \times 10^6$ cells per 15 cm plate. On day 2, cells were either transfected with LysoTag (TMEM192-2xFLAG) using FuGENE 6 or infected with P2 viral stocks expressing the same construct. On day 4, cells were harvested, washed twice with PBS, and scraped in 1 mL of KPBS buffer followed by centrifugation at 1,000 x *g* for 2 min at 4°C. Pelleted cells were resuspended in 1 mL KPBS, and 25 μL was reserved for whole-cell fraction analysis. The remaining 950 μL was gently homogenized with 20 strokes using a 2 mL Dounce homogenizer. The homogenate was centrifuged at 1,000 x *g* for 2 min at 4°C, and 750 μL supernatant was collected. A total of 60 μL was used to determine protein concentration by BCA assay, and 690 μL was incubated with 100 μL of KPBS-prewashed 50% anti-FLAG M2 beads suspension (Sigma) on a gentle rotator for 10 min. Immunoprecipitants were washed three times with KPBS using a DynaMag-2 magnetic rack and eluted with protein loading buffer for further analysis.

For the whole-cell fraction, cells were lysed in RIPA buffer supplemented with a protease inhibitor cocktail (Roche), 100U/mL Benzonase Nuclease (Sigma) at 4°C for 20 min. Cell debris was removed by high-speed centrifugation at 16,100 x *g* for 10 min at 4°C, and the supernatant was collected. Protein samples were separated on 4-12% gradient SDS-PAGE gels and analyzed by western blotting with the indicated antibodies.

Immunoblotting was performed for TMEM192, NPC1, mTOR, LAMTOR1, CNX (Calnexin), GIANTIN, and ERGIC53 in whole-cell lysates and purified lysosomes. CNX, GIANTIN, and ERGIC53 served as negative control, while NPC1 was used as positive control. Quantification of mTOR, LAMTOR1 and NPC1 protein levels in KO and WT cells was performed using ImageJ. This experiment was repeated twice and the results were similar.

**Sterol depletion assays**

Sterol depletion assays were performed as described previously (42). Briefly, on Day 0, $1.5 \times 10^5$ HEK293A WT cells or $2.2 \times 10^5$ TM6SF1 KO cells were seeded on a 6-well plate. On Day 1, the pEG BacMam vector encoding different variants of TM6SF1 or empty vector (EV) was transfected into 6-well plates. After 48 hours, cells were treated with DMEM-high glucose containing 0.5% Lipoprotein Deficient Serum (LPDS) with 1% (w/v) HPCD (Trappsol), plus 10 μM sodium compactin and 50 μM sodium mevalonate at 37°C for 1.5 hour. The cells were then washed with PBS and further cultured in DMEM-high glucose containing 5% LPDS, 10 μM sodium compactin, plus 50 μM sodium mevalonate with or without 50 μM Methyl-β-cyclodextrin (MCD)-cholesterol or DMEM-high glucose containing 0.5% LPDS, 10 μM sodium compactin, plus 50 μM sodium mevalonate with or without 50 μg/mL LDL at 37°C for an additional 2 h. Then, cells were harvested for immunoblot analysis. This experiment was repeated twice and the results were similar.

**Construction of model**

Two all-atom protein structures, (1) protein without cholesterol, (2) protein with cholesterol, were modeled from the atomic coordinates of 1OUP obtained at 2.9-Å resolution. Hydrogen atoms were added using H-build from CHARMM (47a2) (55). The pKa values of all titratable residues were determined at pH 5.5 using the Poisson-Boltzmann Electrostatic Question (PBEQ) Solver in CHARMM-GUI(56).

For each model, the protein structure was modeled in a lipid bilayer consisting of 80% 1-Palmitoyl-2-oleoyl-D-glycero-1-phosphatidylcholine (POPC), 10% 2,3-dioleoyl-D-glycero-1-phosphatidylglycerol (DOPG), and 10% cholesterol. The transmembrane region of the protein was positioned in the lipid bilayer using the CHARMM-GUI (56) and OPM database (57). Next, each protein-membrane system was solvated with explicit TIP3 water (58) and $K^+$ and $Cl^-$ ions corresponding to 0.15 M KCl. Each solvated protein model, consisting of ~161,000 atoms, was simulated in a rectangular box of dimension 125 Å x 125 Å x 149 Å.

**Geometry optimizations and molecular dynamics**

The initial geometry of each solvated protein-membrane complex was optimized using two rounds of 100 steps of steepest descent followed by 100 steps of adopted basis Newton-Raphson (ABNR) to remove any close contacts. All energy minimizations used the all-atom CHARMM36 parameters for proteins, lipids, and ions (59) and the TIP3P model for water molecules (58). Next, the solvated protein-membrane complex was simulated with molecular dynamics (MD) at 310 K according to the following protocol with NAMD (3.0b4) (60): 1) equilibration MD with Langevin dynamics (time step of 1 fs) for 375 ps followed by Langevin dynamics (time step 2 fs) for 1.5 ns; 2) production MD with Langevin dynamics (time step 2 fs) for 200 ns. Periodic boundary conditions were applied to simulate a continuous system. Electrostatic interactions were summed with the Particle Mesh Ewald method (61) grid spacing ~0.87-1.25 Å; fftx 144, ffty 144, fftz 120); a nonbonded cutoff of 12.0 Å was used. Pressure was controlled using a Langevin piston (1.101325 bar), and temperature was controlled with Langevin dynamics with damping coefficient 1.0 $ps^{-1}$. Visualization and data analysis were performed with VMD (version 1.9.4a57) (62) and MD analysis (63).

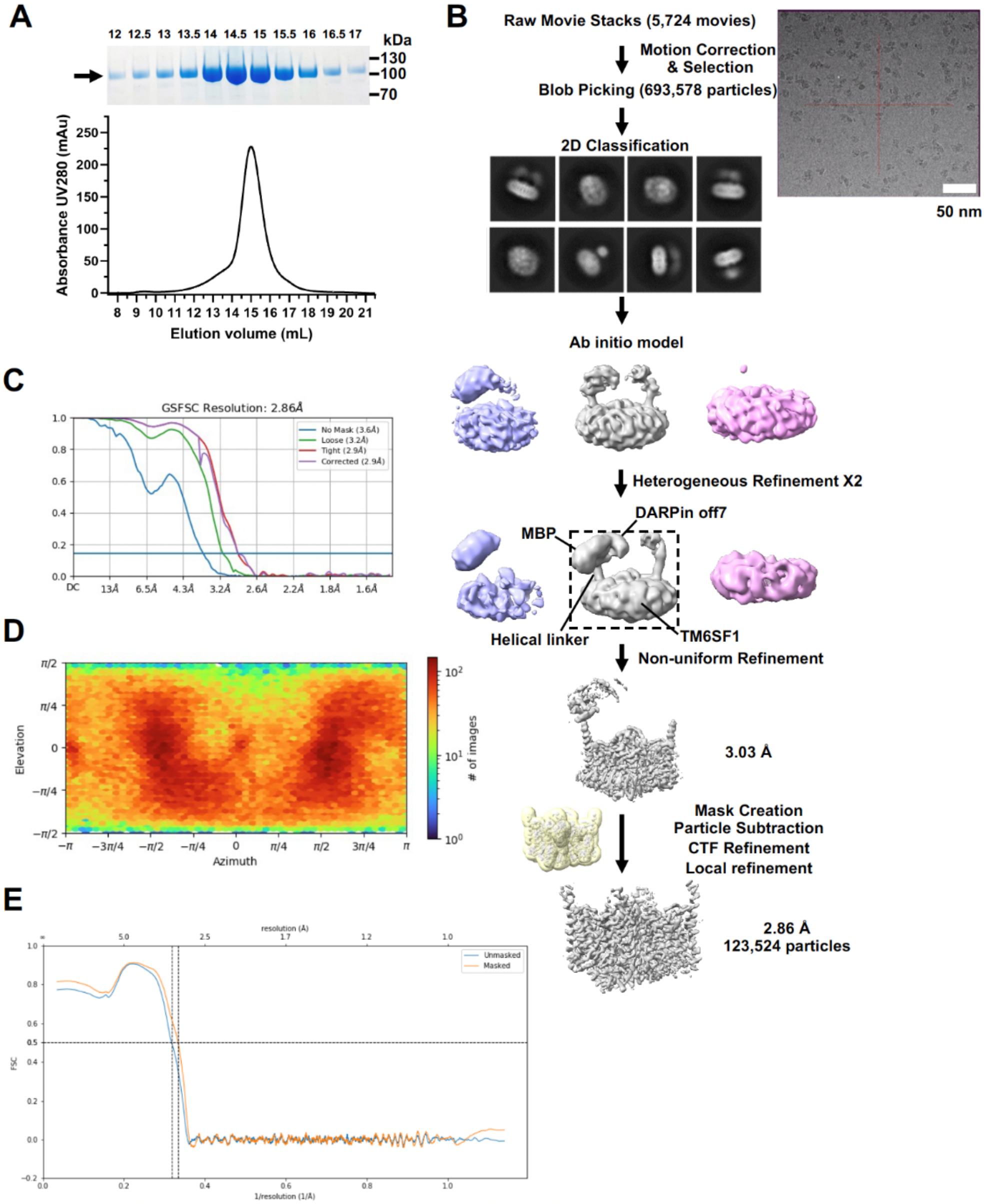

**Fig. S1. Cryo-EM analysis of human TM6SF1$^{MBP}$.** (*A*) A representative size-exclusion chromatogram of TM6SF1$^{MBP}$ protein using a Superose 6 Increase 10/300 GL column in buffer D is shown. The corresponding Coomassie stain of an SDS-PAGE gel with molecular weight markers is displayed on the top. The TM6SF1$^{MBP}$ band is indicated by an arrow. (***B***) Workflow summary of the image processing for TM6SF1$^{MBP}$. (***C***) Fourier shell correlation (FSC) curves between two half maps of TM6SF1 $^{MBP}$. (***D***) Angular distribution of particles used in the final 3D reconstruction of TM6SF1$^{MBP}$. (***E***) FSC curve of the TM6SF1 model to cryo-EM map.

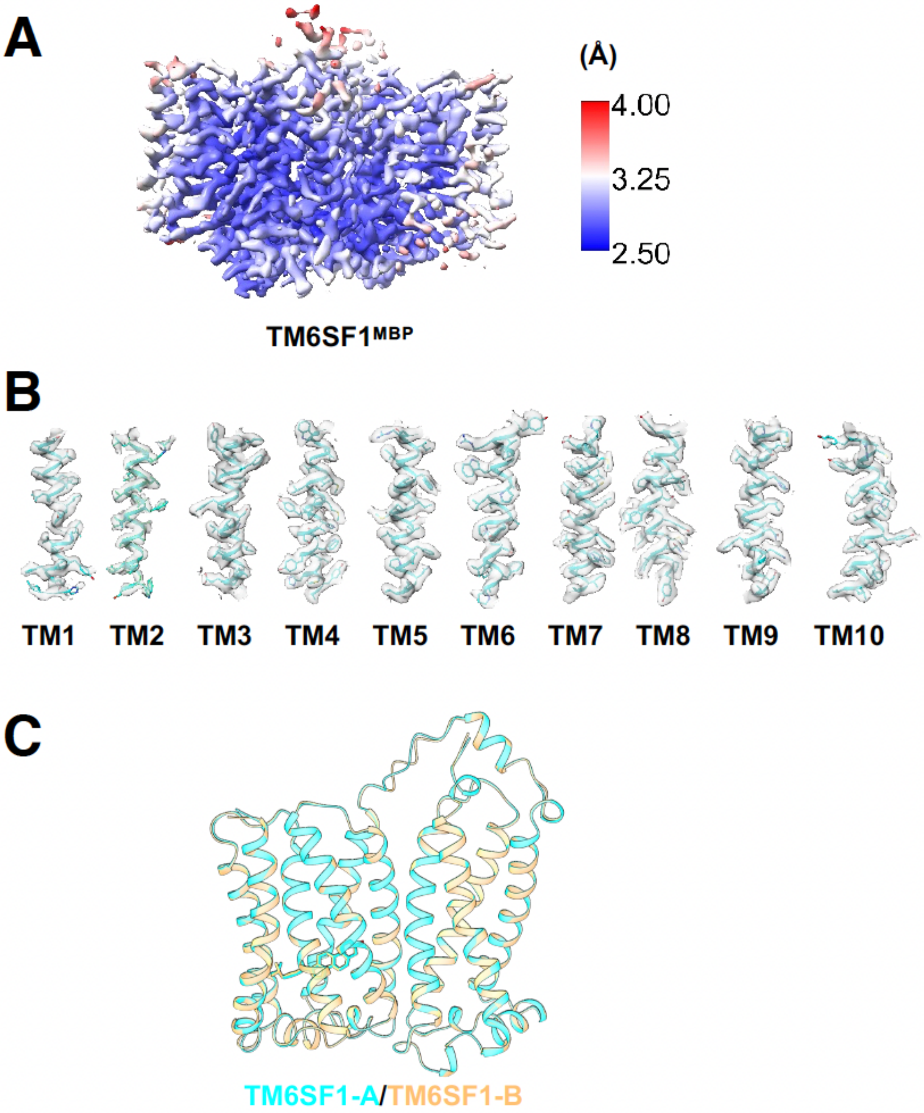

**Fig. S2. Representative cryo-EM density maps of TM6SF1$^{MBP}$.** (***A***) Local resolution map of TM6SF1$^{MBP}$, colored according to local resolution estimates generated using cryoSPARC. The color gradient (blue-white-red) represents a resolution range from 2.5 Å to 4.0 Å. (***B***) Cryo-EM density of the transmembrane (TM) helices of TM6SF1$^{MBP}$. (***C***) The two protomers in the dimer are superimposed and are nearly identical.

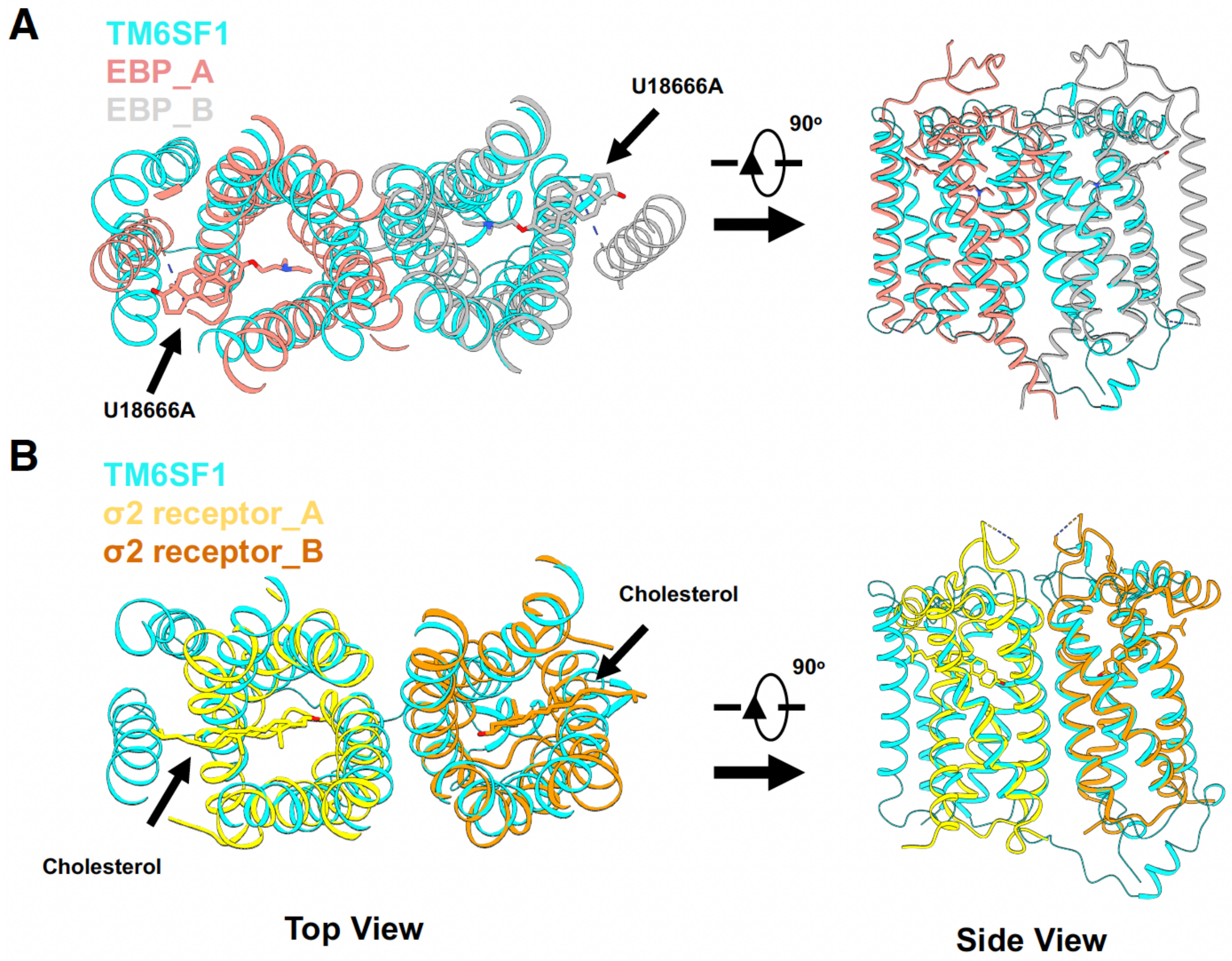

**Fig. S3. Structural comparison of TM6SF1 with EBP and σ2 receptor. (*A*)** Superposition of TM6SF1 (cyan) with EBP. Protomer A and B are pink and gray, respectively (PDB: 6OHT). The sterol-like inhibitor U18666A bound to EBP is shown as a stick. Left: top view; right: side view of the transmembrane helical bundle. **(*B*)** Structural alignment of TM6SF1 (cyan) with the σ2 receptor (protomer A in yellow, protomer B in orange) (PDB: 7MFI). Cholesterol molecules bound to the σ2 receptor are indicated by stick representations. Both top and side views are shown to highlight cholesterol-binding sites.

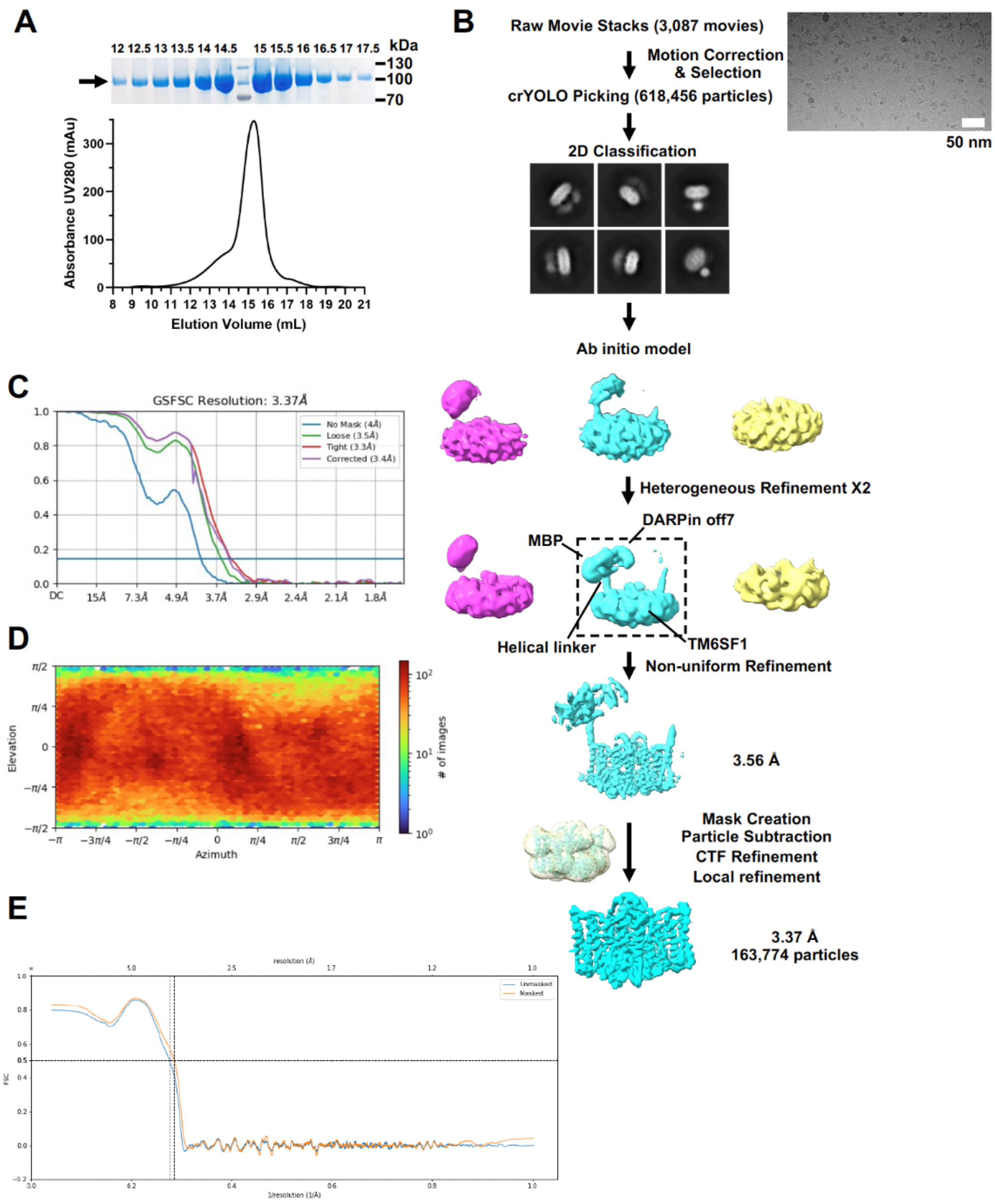

**Fig. S4. Cryo-EM analysis of human TM6SF1$^{\mathrm{MBP/3A}}$.** (***A***) A representative size-exclusion chromatogram of TM6SF1$^{\mathrm{MBP/3A}}$ protein using a Superose 6 Increase 10/300 GL column in buffer D is shown. The corresponding Coomassie stain of an SDS-PAGE gel with molecular weight markers is displayed on the top. The TM6SF1$^{\mathrm{MBP/3A}}$ band is indicated by an arrow. (***B***) Workflow summary of the image processing for TM6SF1$^{\mathrm{MBP/3A}}$. (***C***) Fourier shell correlation (FSC) curves between two half maps of TM6SF1$^{\mathrm{MBP/3A}}$. (***D***) Angular distribution of particles used in the final 3D reconstruction of TM6SF1$^{\mathrm{MBP/3A}}$. (***E***) FSC curve of the TM6SF1$^{\mathrm{3A}}$ model to cryo-EM map.

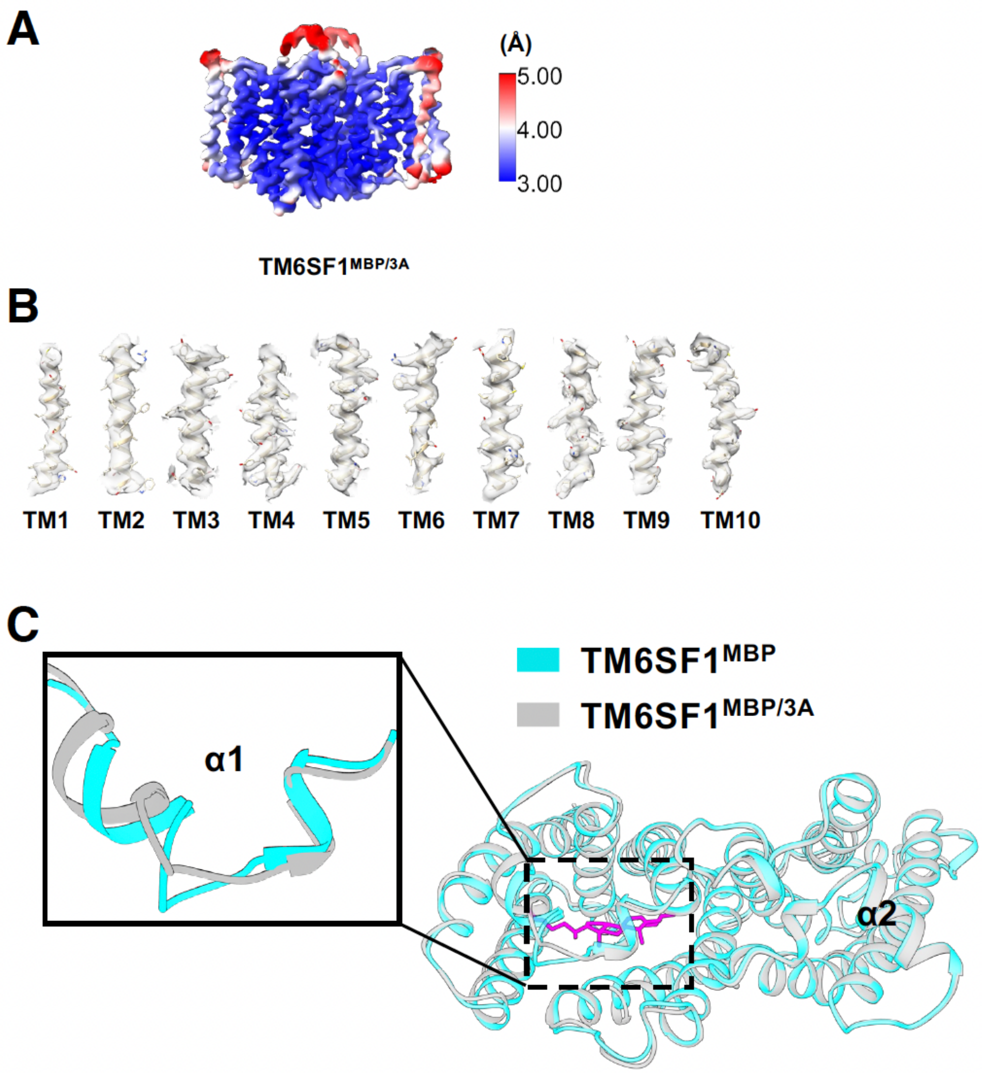

**Fig. S5. Representative cryo-EM density maps of TM6SF1$^{MBP/3A}$. (*A*)** Local resolution map of TM6SF1$^{MBP/3A}$, colored according to local resolution estimates generated using cryoSPARC. The color gradient (blue-white-red) represents a resolution range from 3 Å to 5 Å. **(*B*)** Cryo-EM density of the transmembrane helices of TM6SF1$^{MBP/3A}$. **(*C*)** Structural superposition of TM6SF1$^{MBP}$ (cyan) and TM6SF1$^{MBP/3A}$ (grey) reveal conformational changes of the cholesterol-binding pocket. The cholesterol molecule is shown as sticks (magenta). A zoomed-in view highlights structural rearrangements in the α1 helix of the mutant protein which are likely to affect cholesterol binding.

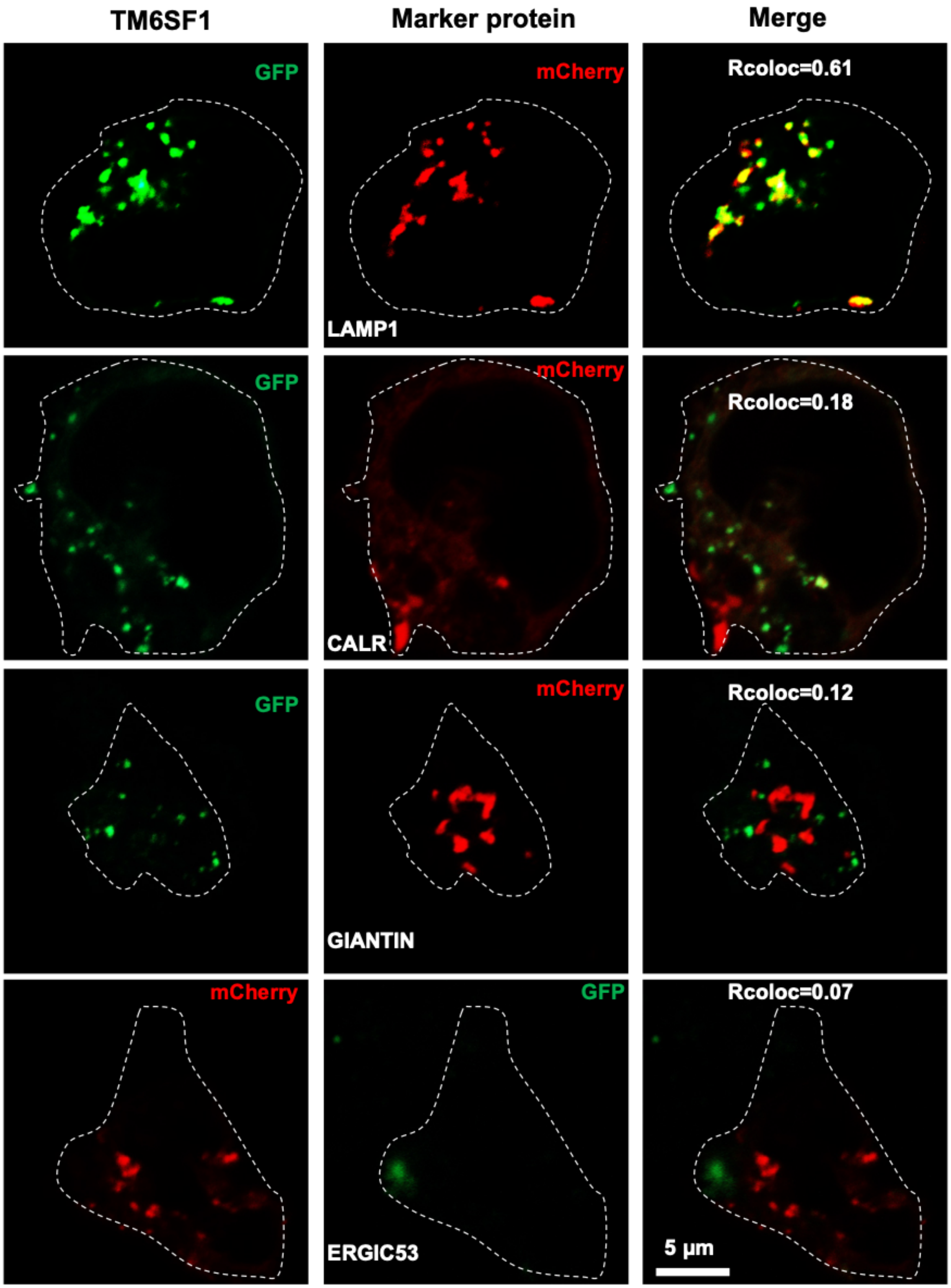
TM6SF1
Marker protein
Merge
GFP
mCherry
Rcoloc=0.61
LAMP1
GFP
mCherry
Rcoloc=0.18
CALR
GFP
mCherry
Rcoloc=0.12
GIANTIN
mCherry
GFP
Rcoloc=0.07
ERGIC53
5 μm

394

**Fig. S6. Subcellular localization of TM6SF1.** TM6SF1 localizes to lysosomes. On day 0, human HEK293A cells were seeded on 22 mm glass coverslips pre-coated with poly-D-lysine in medium B supplemented with 10% FCS at a density of $2.5 \times 10^5$ cells per well in a 6-well plate. On day 1, cells were transfected using FuGENE6 with the following plasmid combinations: GFP-tagged TM6SF1 and mCherry-tagged LAMP1 (lysosomal marker), GFP-tagged TM6SF1 and mCherry-tagged CALR (ER marker), GFP-tagged TM6SF1 and mCherry-tagged GIANTIN (Golgi marker), mCherry-tagged TM6SF1 and GFP-tagged ERGIC53 (ERGIC marker). On day 2, cells were fixed and analyzed by fluorescence microscopy. Colocalization was quantified using Pearson's correlation coefficient (Rcoloc), and the values represent the mean analyzed in three independent experiments. Scalebar, 5 µm.

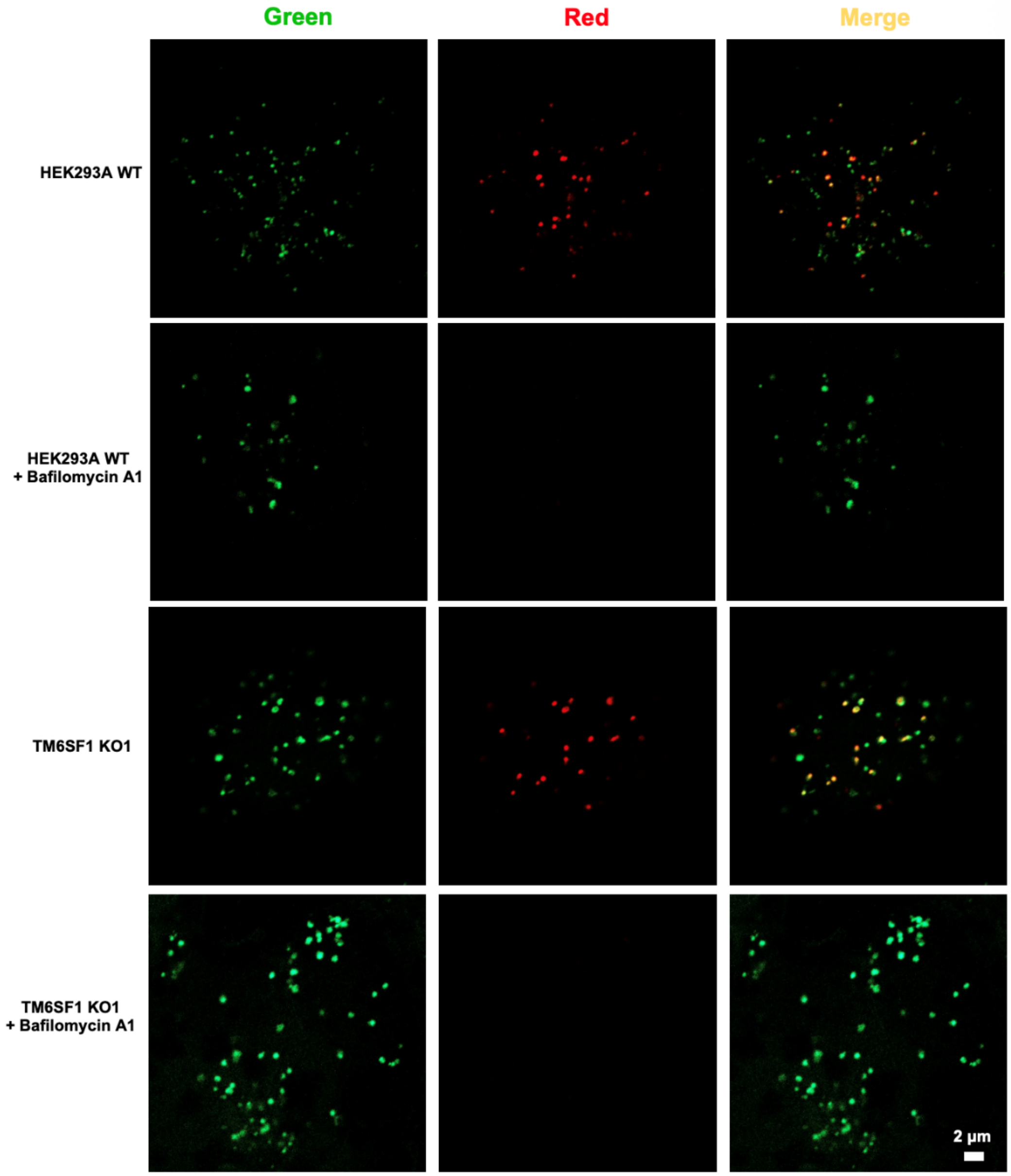

**Fig. S7. No effect of inactivation of TM6SF1 on lysosomal acidification**. On day 0, WT HEK293A cells and TM6SF1 KO cells were set up in medium B supplemented with 10% FCS at a density of 2.5 × $10^5$ cells per well. On day 1, cells were incubated with 200 µL of LysoPrime Green at 37 °C. After 30 min, cells were washed and incubated with 200 µL of pHLys Red +/- bafilomycin A1 at 37°C for 30 min. The cells were then washed and maintained in medium B supplemented with 10% FCS. The stained cells were visualized under a confocal microscope. Treatment of cells with bafilomycin A1 was used as a control for the experiment.

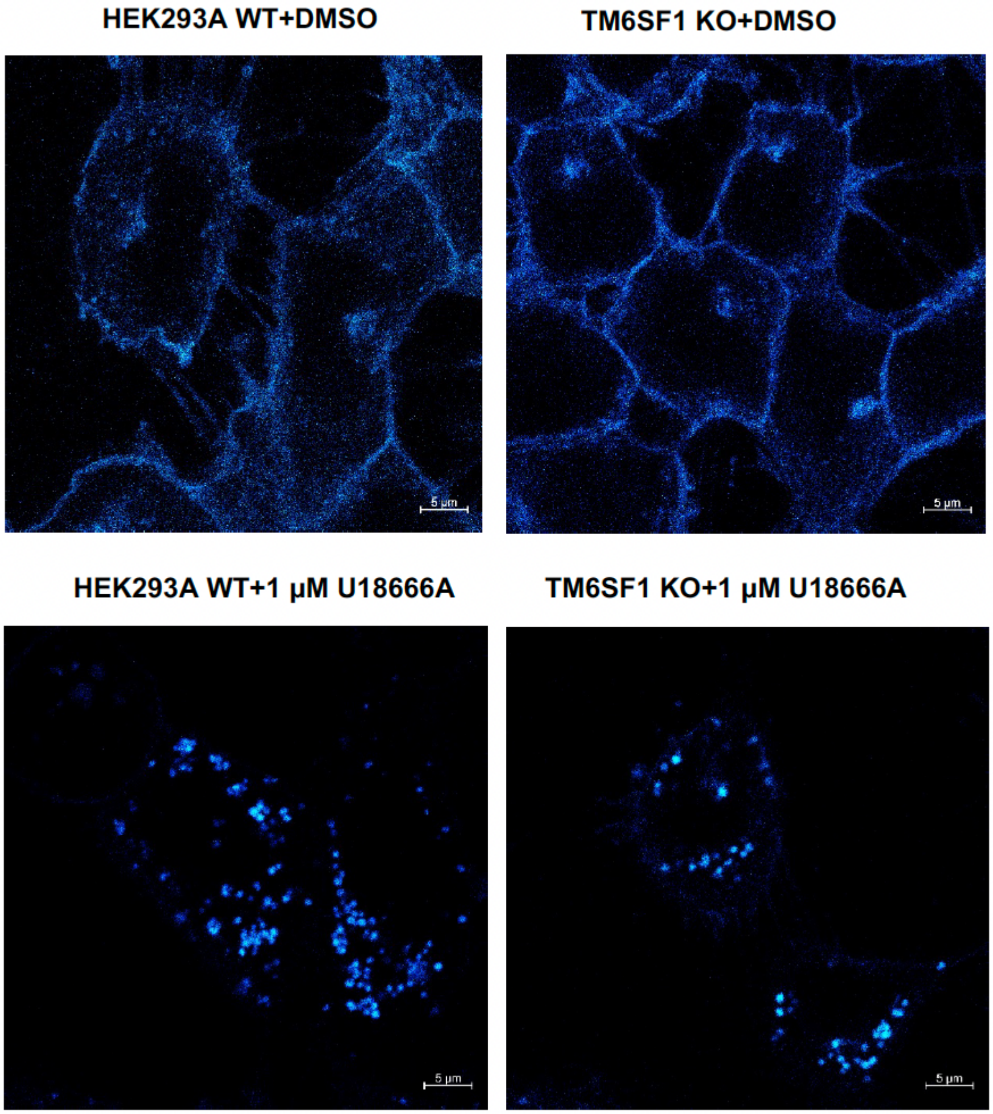

**Fig. S8. Cholesterol distribution in WT HEK293A cells and TM6SF1 KO cells.** On day 0, WT and TM6SF1 KO cells were seeded onto 22 mm glass coverslips in medium C supplemented with 5% FCS. On day 1, cells were treated with DMSO (solvent control) or 1 µM U18666A. On day 2, cells were switched to cholesterol-depletion medium C. After 12 h of incubation, cells were switched to medium C supplemented with 30 µM compactin, 200 µM mevalonate, and 10 % FCS. After 6 h, cells were then fixed and stained with filipin to visualize free cholesterol. Scalebar, 5 µm.

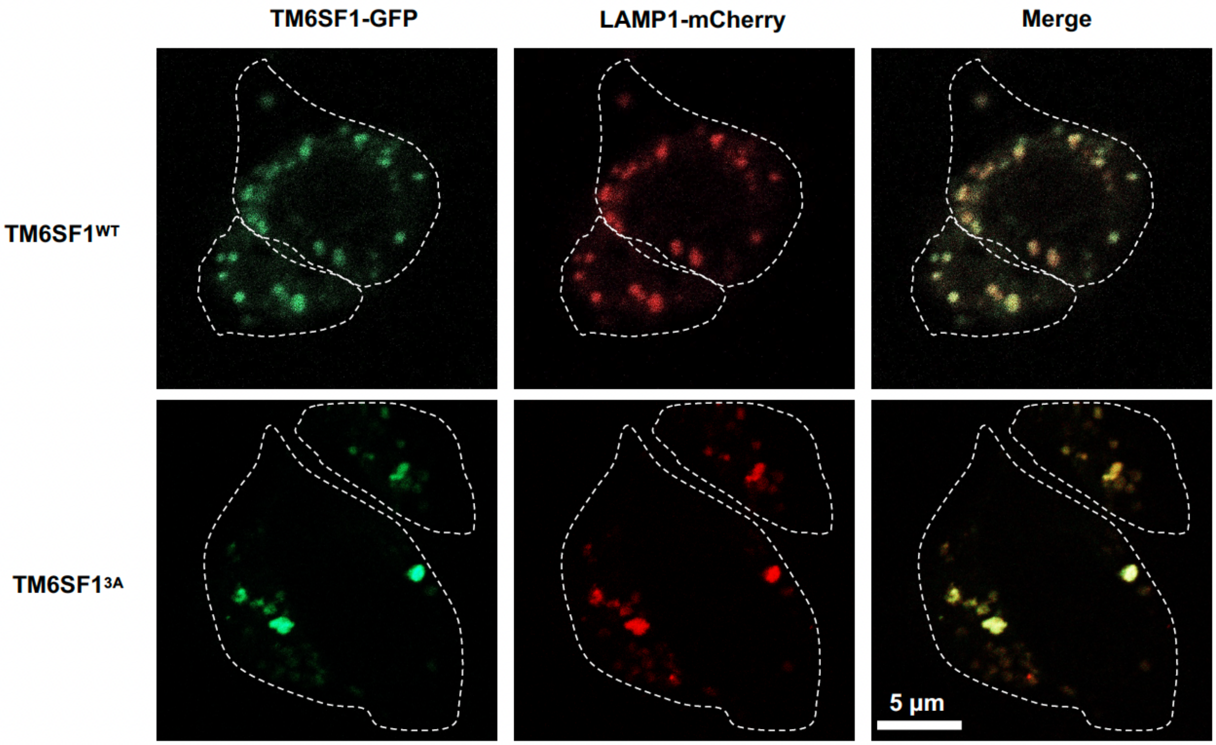

**Fig. S9. Subcellular localization of TM6SF1$^{3A}$.** On day 0, TM6SF1 KO cells were seeded on 22 mm glass coverslips pre-coated with poly-D-lysine in medium B supplemented with 10% FCS at a density of 2.5 × 10 5 cells per well. On day 1, cells were transfected with the following plasmid combinations: GFP-tagged TM6SF1 and mCherry-tagged LAMP1, and GFP-tagged TM6SF1$^{3A}$ and mCherry-tagged LAMP1. On day 2, cells were fixed and analyzed by fluorescence microscopy. Scalebar, 5 µm.

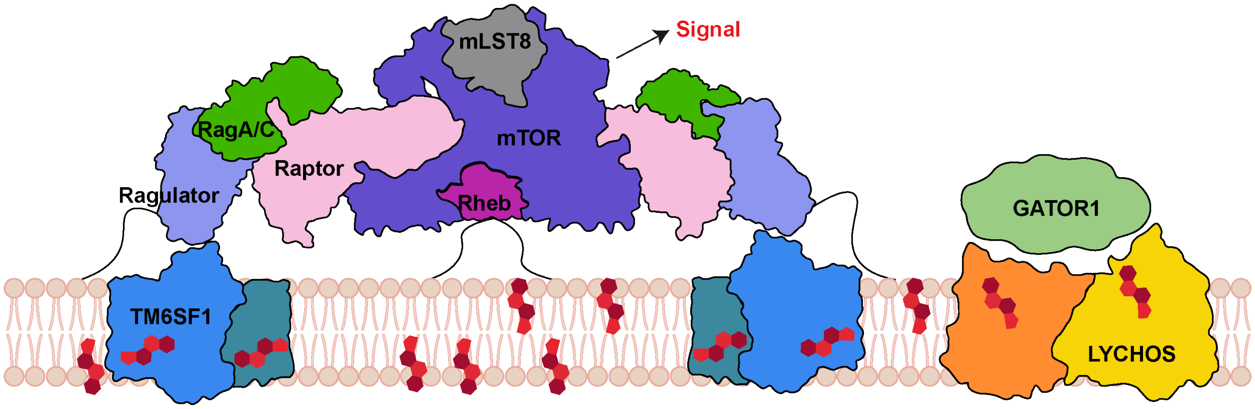

**Fig. S10. Proposed model of mTORC1 docking mediated by cholesterol bound TM6SF1.** Cholesterol is colored in red.

**Table S1 Cryo-EM data collection, refinement and validation statistics.**

| **Structure** | **TM6SF1$^{MBP}$** | **TM6SF1$^{MBP/3A}$** |
|---|---|---|
| **EMDB** | **EMD-75471** | **EMD-75606** |
| **PDB** | **10UP** | **11BR** |
| **Data collection and processing** | | |
| Magnification | 165,000 | 105,000 |
| Voltage (kV) | 300 | 300 |
| Pixel size (Å) | 0.738 | 0.834 |
| Defocus range (μm) | 0.8-1.8 | 0.8-1.8 |
| Electron exposure ($e^-$/ Å$^2$) | 60 | 60 |
| Symmetry imposed | C1 | C1 |
| Initial particles (No.) | 693,578 | 618,456 |
| Final particle images (No.) | 123,524 | 163,774 |
| Map resolution (Å) | 2.86 | 3.37 |
| FSC threshold | 0.143 | 0.143 |
| Map sharpening *B* factor (Å$^2$) | -86.5 | -127.6 |
| **Refinement** | | |
| Initial model used (PDB code) | AlphaFold2 | 10UP |
| Model Resolution (Å) | 3.0 | 3.5 |
| FSC threshold | 0.5 | 0.5 |
| Model Composition | | |
| Chains | 2 | 2 |
| Non-hydrogen atoms | 5824 | 5746 |
| Protein residues | 726 | 726 |
| Ligands | 2 | 0 |
| Water | 0 | 0 |
| *B* factors (Å$^2$) | | |
| Protein | 66.23 | 140.36 |
| Ligand | 68.73 | N/A |
| Water | N/A | N/A |
| R.M.S. deviations | | |
| Bond lengths (Å) | 0.002 | 0.003 |
| Bond angles (°) | 0.497 | 0.528 |
| Validation | | |
| MolProbity score | 0.96 | 0.99 |
| Clashscore | 1.62 | 2.16 |
| Rotamers outliers (%) | 1.15 | 0.66 |
| Ramachandran plot (%) | | |
| Outliers | 0.00 | 0.00 |
| Allowed | 1.66 | 1.94 |
| Favored | 98.34 | 98.06 |
